# Combined analysis of the time-resolved transcriptome and proteome of plant pathogen *Xanthomonas oryzae* pv. *oryzae*

**DOI:** 10.1101/2020.10.02.324541

**Authors:** Seunghwan Kim, Wooyoung Eric Jang, Min-Sik Kim, Jeong-Gu Kim, Lin-Woo Kang

## Abstract

*Xanthomonas oryzae* pv. *oryzae* (Xoo) is a plant pathogen responsible for causing bacterial blight in rice. The immediate alterations in Xoo upon initial contact with rice are essential for pathogenesis. We studied time-resolved genome-wide gene expression in pathogenicity-activated Xoo cells at the transcriptome and proteome levels. The early response genes of Xoo include genes related to cell motility, inorganic ion transport, and effectors. The alteration of gene expression is initiated as early as few minutes after the initial interaction and changes with time. The time-resolved comparison of the transcriptome and proteome shows the differences between transcriptional and translational expression peaks in many genes, although the overall expression pattern of mRNAs and proteins is conserved. The discrepancy suggests an important role of translational regulation in Xoo at the early stages of pathogenesis. The gene expression analysis using time-resolved transcriptome and proteome provides unprecedented valuable information regarding Xoo pathogenesis.

## Introduction

Rice (*Oryza sativa* L.) is the most widely consumed staple food, sustaining two-thirds of the world’s population (Jackson, 2016). Bacterial blight of rice caused by *Xanthomonas oryzae* pv. *oryzae* (Xoo) is a devastating disease for which an effective pesticide has not been developed yet; it is known to cause severe yield losses of up to 50% in several rice-growing countries (Oliva et al., 2019). The demand for rice is expected to increase by at least 25% by 2030, owing to the rapidly growing world population, environmental stress arising in response to climate change, and pathogen pressure (Li, Wang, & Zeigler, 2014). Further, the Green Revolution has resulted in a shift in rice cultivation, from varied traditional landraces to limited high-yielding varieties, via artificial selection. This has resulted in the co-evolution of crop pathogens including Xoo with the selected host races in the modern agricultural ecosystem (Quibod et al., 2020).

The pathogen-host system of Xoo and rice serves as an ideal agricultural model to study crop diseases in a field setting at the molecular level; this is facilitated by the early elucidation of the whole genome structure of both Xoo and rice (Jackson, 2016; Lee et al., 2005). Xoo typically invades rice leaves through the wounds or hydathodes and replicates in the xylem vessels to cause disease (Mew, Alvarez, Leach, & Swings, 1993). Rice contains a two-tiered innate immune system, consisting of pathogen-associated molecular pattern- and effector-triggered immunity, which protects against Xoo and initiates the hypersensitivity response at the infection site (Jones & Dangl, 2006). In Xoo-rice interactions, Xoo injects effectors into rice cells to modulate the cellular activities of the host to promote pathogenesis (Tsuge, Furutani, & Ikawa, 2014). The early interactions between Xoo and rice at the infection site determine the fate of infection, i.e., occurrence of disease or initiation of the immune response. The environmental conditions prevalent at the site of infection are varied and complex, and our understanding of alterations in the Xoo cells in response to the initial interactions with rice is limited.

Transcription and translation are tightly coupled in bacteria and can occur simultaneously in the cytosol (Hershey, Sonenberg, & Mathews, 2019). Although proteins are the final functional products of genes, the quantity of specific mRNA molecules often represents the expression level of a gene at a given time point with the well-established RNA sequencing (RNA-Seq) technology. In comparison, high resolution mass spectrometry-based quantitative proteomics is a more recent analytical technique, and still has a lower coverage of protein products, requires greater sample quantity, and is more expensive (Schubert, Rost, Collins, Rosenberger, & Aebersold, 2017).

We had previously developed an *in vitro* pathogenicity assay to recapitulate Xoo-rice interactions at the site of infection by treating Xoo cells with the rice homogenate (RLX), and assessed the time-resolved changes in the transcriptome (S. Kim et al., 2016; S. H. Kim et al., 2011; S. Kim et al., 2013). The *in vitro* pathogenicity assay provides high signal to noise data with Xoo cells synchronized with respect to the timing of pathogenicity activation. Transcriptome data from RNA-Seq experiments revealed that most virulence genes of Xoo were upregulated within an hour of the initial interaction with RLX, and these upregulated genes were related to bacterial motility, inorganic ion transport, hypersensitive response and pathogenicity (*hrp*), bacterial toxins and effectors of avirulence (*avr*), plant cell wall degradation, and extracellular polysaccharide synthesis and secretion (S. Kim et al., 2016).

Here, we expand the study of gene expression in pathogenicity-activated (P-activated) Xoo from the transcriptome to the proteome. The time-dependent expression of specific mRNAs and proteins represents the kinetics of transcriptional and translational expression of certain genes and allow the visualization of the immediate alterations in gene expression in P-activated Xoo.

## Results

### Time-resolved proteome data

We coupled LC-MS/MS technology with an *in vitro* assay system to obtain the time-resolved proteome data for P-activated Xoo cells (Scheme 1). The *in vitro* assay system recapitulated the initial interaction between Xoo cells and damaged rice leaf tissues at the site of infection; this was achieved by adding fresh RLX—prepared by grinding the leaves of a Xoo-susceptible rice cultivar (Milyang 23) in liquid nitrogen—to a Xoo cell culture in the mid-exponential phase. Samples for proteome analysis were collected from RLX-treated (P-activated) and untreated (control) Xoo cells at 0, 30, 60, 90, and 120 min after RLX treatment (Table S1).

**Scheme 1.**
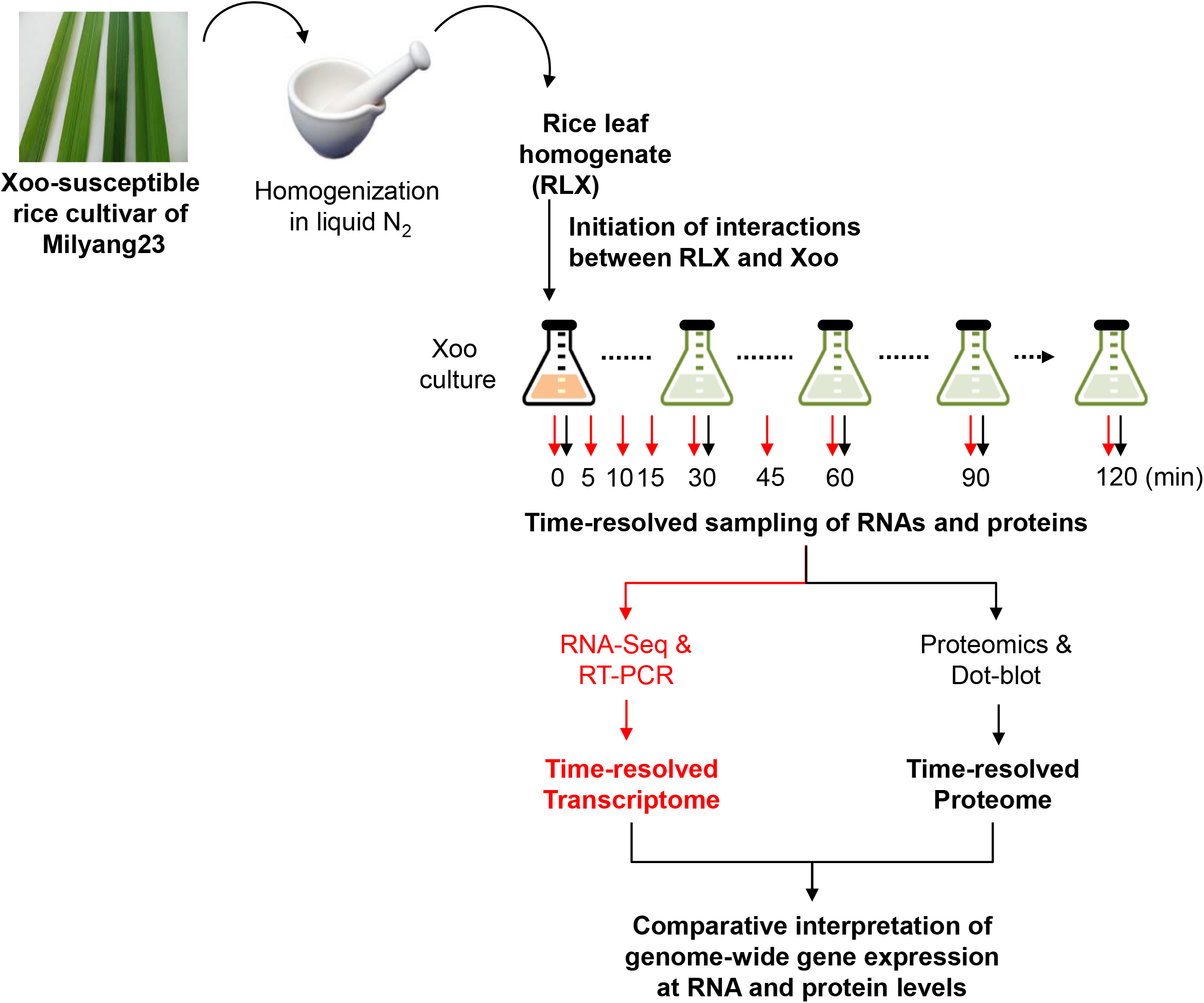
Schematic representation of the in vitro assay system and combined analysis of the time-resolved transcriptome and proteome using RNA-Seq and LC-MS/MS

Analysis on UniProt revealed that the total 4,956 predicted open reading frames in the Xoo genome (KACC10331) corresponded to 4,382 proteins in the proteome, of which 2,589 proteins (59%) were identified for at least one time point and 2,296 proteins (52%) were detected in both replicates (Figure S1A). Median sequence coverages for total identified proteins were 24% and 23% for each of the independent duplicate experiments (Figure S1B). A total of 20,963 and 19,684 non-redundant peptides were identified in both replicates, with 47,025 and 41,428 peptide-spectrum matches, respectively (Figure S1C).

Protein abundance values obtained after quantile-normalization (Figure S2A) were used for pairwise comparisons; the Pearson’s correlation coefficients corresponding to the abundance values showed close correlations (0.98-0.99), indicating comparable cellular concentration of most proteins (Figure S2B). The smallest correlations were observed for the RLX-treated samples at 30 min, indicating greater changes in the proteome during the initial 30 min; this was consistent with the transcriptome data. We further performed a principal component analysis to determine the relationships between the assessed samples. Figure S2C shows that the P-activated sample at 0 min clustered closely with all control samples, whereas the P-activated samples at other time points were more spread out. The proteome of the P-activated Xoo cells at 30 min was considerably different from that of P-activated Xoo at other time points; this was consistent with the results of the pairwise multi-scatter plot (Figure S2B).

Time-supervised hierarchical clustering was performed to determine temporal and synchronized changes in protein abundance. Two distinct synchronized patterns—one decreasing and the other increasing in response to RLX treatment—were observed (Figure 1). The two lists including proteins exhibiting the two patterns were used for STRING analysis, which produced two interaction networks (Figure S3A-B). Several genes related to cell motility, inorganic ion transport, and transcriptional regulators were immediately responsive to the pathogenicity signal (Table S2); these have been described in detail in the discussion section.

**Figure 1.**
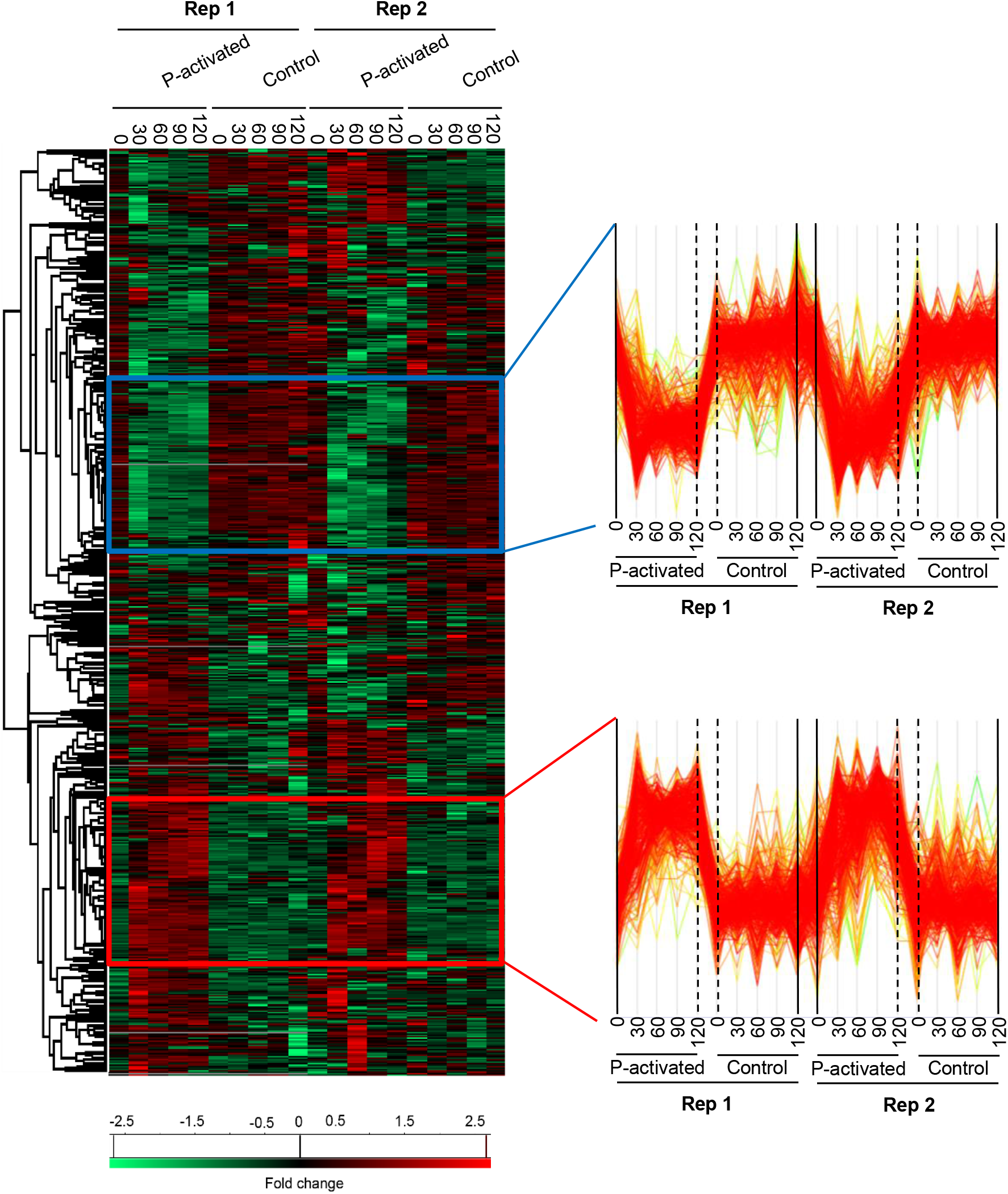
Time-supervised hierarchical clustering of duplicate pathogenicity-activated proteome datasets. The heatmap (left) of genes clustered based on the similarities in the protein expression pattern over time is presented for duplicate samples of pathogenicity-activated and control Xoo cells. Low to high expression is indicated by a change in color from green to red. Selected cluster profile patterns (right) are presented for the downregulated (blue box) and upregulated (red box) genes in the P-activated Xoo cells.

### Up and down regulated proteins

Quantitative proteome data obtained from P-activated and control Xoo cells at every 30 min allowed the visualization of the three-dimensional protein expression data in terms of the genes, time intervals, and expression levels (Figure 2 and Table S3). The expressed protein level of each gene from P-activated Xoo cells are divided by that of the same gene from control at each time point to calculate the fold change of time-dependent protein expression level of the specific gene. For all the open reading frames, approximately 93 (2.0%), 213 (4.5%), and 468 (9.9%) proteins were upregulated by more than 200% (2-fold), 50%, and 20% at 30 min, respectively, and approximately 7 (0.1%), 93 (2.0%), and 561 (11.9%) proteins were downregulated to less than 25% (2-fold), 50%, and 80% at 30 min, respectively (Table S4).

**Figure 2.**
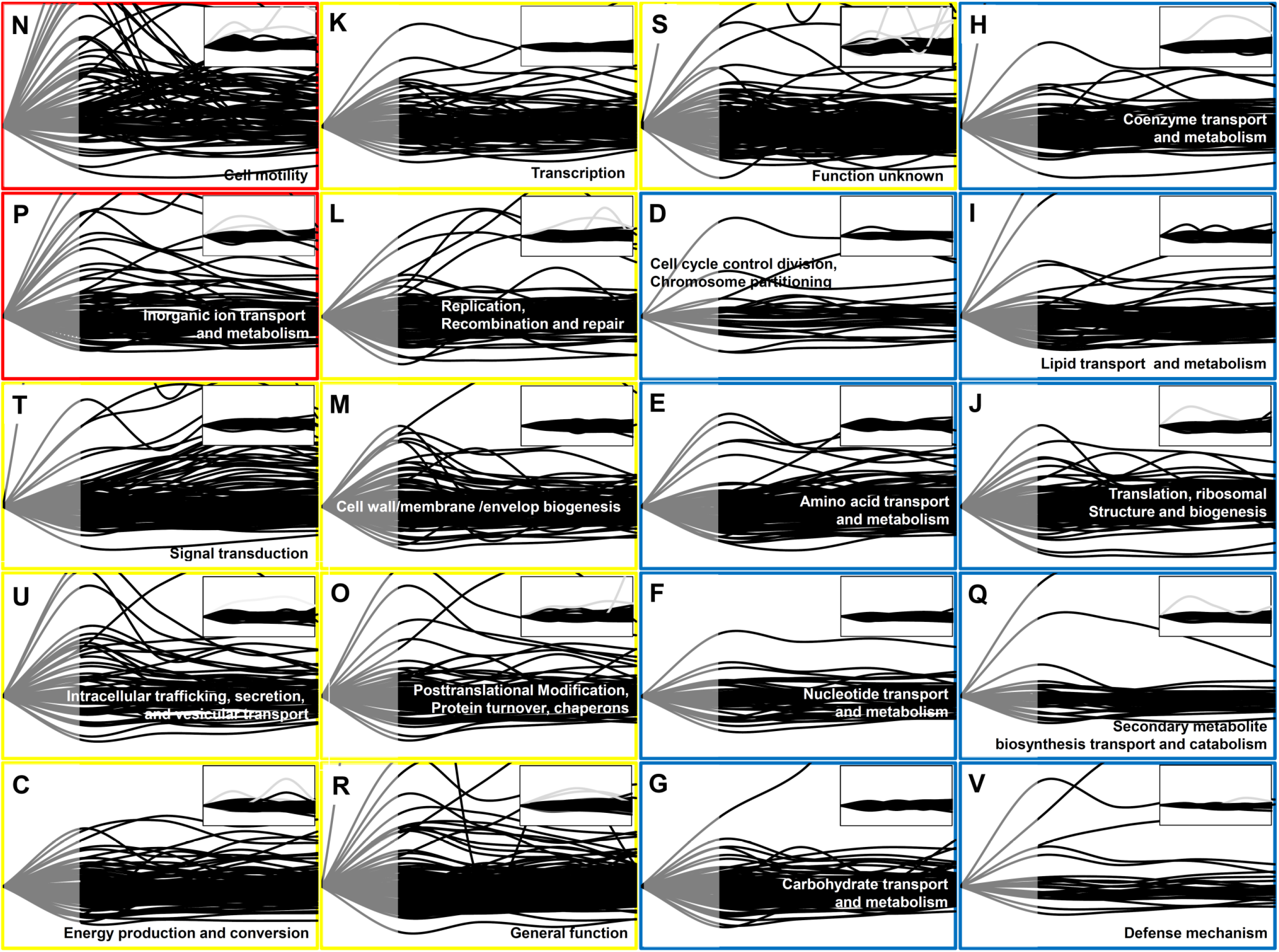
Time-resolved protein expression patterns associated with the COG categories. Two functional categories of genes (COG) associated with the greatest changes in the protein level are indicated by red boxes, nine functional COGs associated with moderate changes are indicated by yellow boxes, and other nine functional COGs associated with minor changes in the protein level are indicated by blue boxes. The inset for each functional COG indicates the control (untreated cells). The Y-axis represents the fold change in the protein expression level in comparison with that at 0 min, and the maximum value on the Y-axis was set as 3. The X-axis indicates the time from 0 to 120 min. Each line indicates the protein expression level of a gene. The protein expression level is represented as grey (from 0 to 30 min) and black (from 30 to 120 min) lines. Note that in contrast to the transcript data, the proteome data was not available between 0 and 30 min (grey part).

In case of the duration of expressed proteins in P-activated Xoo, only 8 (0.2%), 32 (0.7%), and 75 (1.6%) proteins were upregulated for entire 120 min by more than 200% (2-fold), 50%, and 20%, respectively and 0 (0%), 1 (0.02%), and 87 (1.8%) proteins were downregulated for the same 120 min to less than 25% (2-fold), 50%, and 80%, respectively (Table S5), indicating that most proteins were temporarily up- or downregulated in the *in vitro* assay.

The proteins at 30 min presented the highest change at the expression level within 120 min. More than 90 proteins (1.9%) were upregulated by more than 2-fold at 30 min in each dataset, whereas 27 proteins were upregulated in both datasets from the duplicate experiments (Table S6). Almost half of the 27 upregulated proteins were related to cell motility and ion uptake, seven were related to chemotaxis and motility, and five were related to transporters and pumps. At the same 30 min, more than 90 proteins were downregulated by more than 50%, 17 proteins were downregulated in both datasets (Table S7), including transcription-related proteins—such as sigma-54 modulation protein and MetE/MetH family transcriptional regulator—and cell division- and cell cycle-related proteins.

### Clusters of Orthologous Groups of proteins (COGs)

To study the global gene expression pattern based on gene function, we superimposed time-dependent protein expression levels as per the functional categories in the Clusters of Orthologous Groups of proteins (COGs) database (Figure 2), which were grouped into three classes (red, yellow, and blue) depending on the observed pattern. To facilitate the comparison of time-dependent protein expression levels, the expression level at 0 min was set as the reference level (=1) for each gene, and the fold change in the protein expression level was calculated at each time point, as in case of the RNA-Seq data analysis (S. Kim et al., 2016).

The most prominent changes in protein expression in Xoo cells were observed for proteins associated with cell motility (N) and inorganic ion transport and metabolism (P), which were placed in the red class. In category N, two major expression peaks, indicating more than 2-fold up-regulation, were detected at 30 and 90 min. In category P, upregulated proteins peaked at around 30 min. Proteins grouped in the yellow class exhibited moderate changes in their expression level, and were divided into nine functional categories, including signal transduction (T), intracellular trafficking, secretion, and vesicular transport (U), energy production and conversion (C), transcription (K), replication (L), cell wall/membrane/envelop biogenesis (M), and protein turnover (O). The blue class included proteins associated with other nine categories, the expression levels of which exhibited little change when compared with those at 0 min, albeit except for a few proteins.

### Comparison of mRNA and protein levels

We compared mRNA and protein levels of each gene in the transcriptome and proteome (Figure 3; thin dashed red lines of mRNAs and thick dashed and solid black lines of proteins). Both transcriptome and proteome data were obtained for up to 120 min after RLX treatment. The experimental procedure used to prepare protein samples for LS-MS/MS required at least 30 min, whereas that for mRNA samples required at least 5 min; the time interval for the proteome data was set to 30 min, whereas that for transcriptome data was set to 5-30 min. Due to the different sampling times, the transcriptome data included mRNA levels at additional time points of 5, 10, 15, and 45 min, whereas the proteome data did not include protein levels between 0 and 30 min, in which case the protein levels (thick dashed black lines) were extrapolated based on the protein levels at other time points using non-linear regression.

**Figure 3.**
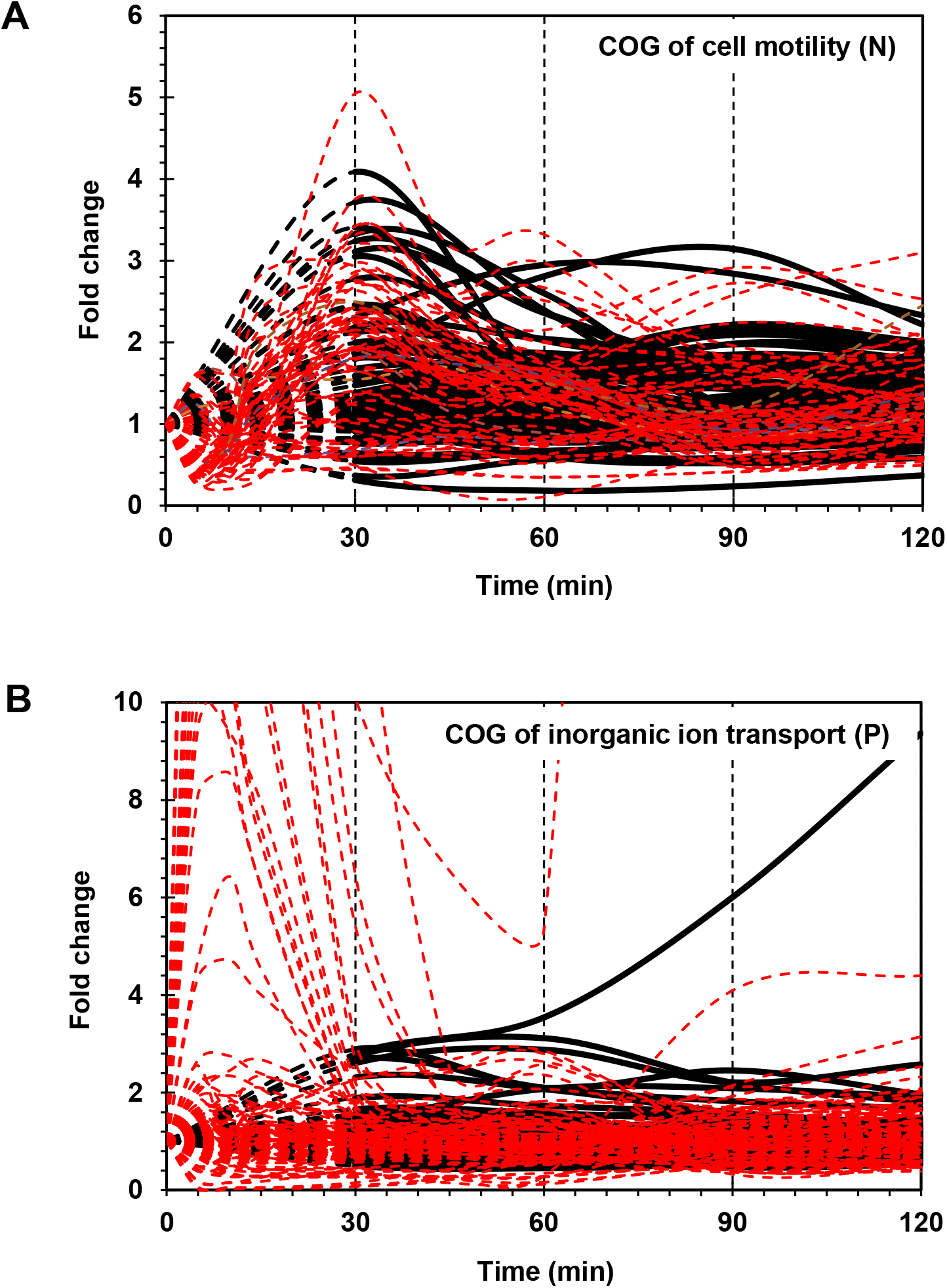
Time-resolved mRNA and protein levels of genes associated with cell motility and inorganic ion transport. (A) Time-resolved expression of cell motility-related genes. The mRNA expression levels from RNA-Seq are indicated by dashed red lines. The protein expression levels from LC-MS/MS are indicated by black dashed (from 0 to 30 min) and black solid (from 30 to 120 min) lines. (B) Time-resolved expression of inorganic ion transport-related genes. The mRNA expression levels from RNA-Seq and protein expression levels from LC-MS/MS are indicated as in (A). The Y-axis represents the fold change.

#### Flagella and chemotaxis-related genes

Flagella and chemotaxis-related genes encode more than 40 proteins, including structural components and assembly factors of flagellar hook-basal body and filament and chemotaxis proteins (Mukherjee & Kearns, 2014). Two dominant hierarchical clusters, with distinct synchronized patterns of increasing and decreasing proteome concentration (indicated by red and blue, respectively), were identified (Figure 1). The red cluster of 359 proteins included 21 (53%) of the 40 genes related to the flagellar assembly pathway, of which 20 genes were included in the red cluster (Table S2).

We grouped the flagella and chemotaxis-related genes into three gene clusters, i.e., groups I-A, I-B, and II, based on their positions in the genome (Figure 4A). Genes in cluster I-A and I-B were associated with flagellar machinery, such as flagellar basal body hook and type III secretion system (T3SS), whereas cluster II included chemotaxis-related genes.

**Figure 4.**
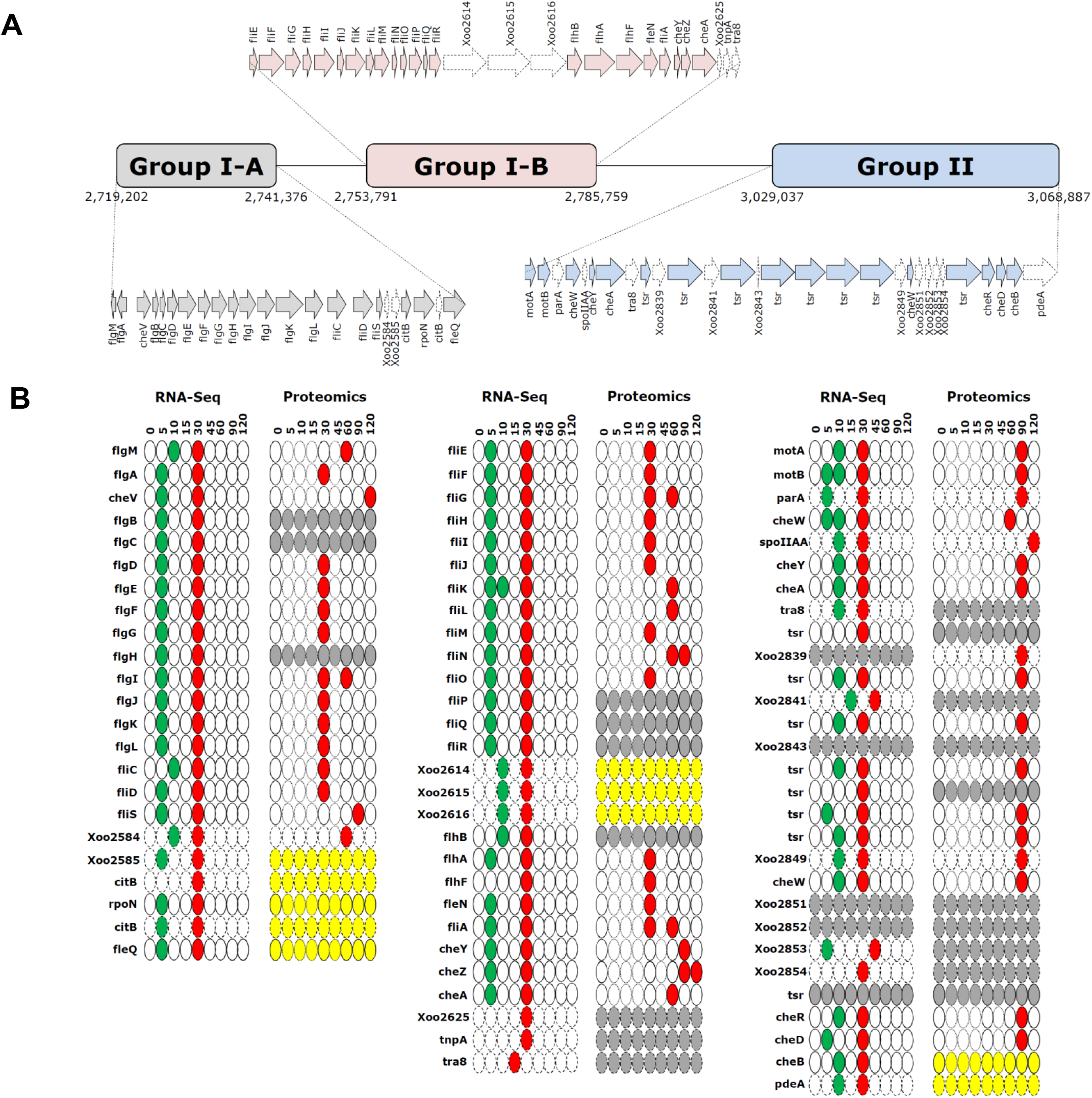
Gene clusters and time-resolved expression patterns of cell motility-related genes. (A) Gene clusters of flagellar biosynthesis-related genes (groups I-A and I-B) and chemotaxis genes (group II). (B) Time-resolved mRNA and protein expression levels of genes in groups I-A, I-B, and II. The down- and upregulation peaks are shown in green and red, respectively. The yellow and grey ovals indicate unaltered and undetected expression levels. The thin-bordered oval (at 5, 10, 15, and 45 min) for the proteome data is to ensure consistency with the pictorial format of RNA-Seq data. The dotted arrows and ovals indicate genes that are not directly related to flagellar biosynthesis and chemotaxis. Time is expressed in min.

Superimposition of transcriptome and proteome data of cell motility related genes revealed clearly superimposed upregulated peaks at 30 min (Figure 3A). The transcriptome data revealed that most flagella and chemotaxis-related genes were regulated in a similar pattern, i.e., genes in all clusters of I-A, I-B, and II were downregulated at 5 min and upregulated at 30 min. However, the proteome data presented a different expression pattern between clusters I-A and I-B and cluster II. Proteins in clusters I-A and I-B were upregulated at 30 min, consistent with the transcriptome data. But proteins in cluster II were upregulated at 90 min, exhibiting a delay of 1 h when compared with the transcriptome data (Figure S4).

#### Inorganic ion transport and metabolism genes

Iron uptake genes play a key role in pathogenicity during the early stages of host-pathogen interactions (Garau et al., 2004). TonB-dependent receptors (TBDRs) are bacterial outer membrane proteins that bind and transport ferric chelates of siderophores. Several annotated TBDR genes have been identified in the Xoo KACC10331 genome, including *IroN, FyuA, FecA, BtuB, FhuA, CirA*, and *FepA* (S. Kim et al., 2016). Of these, *FecA* (*Xoo0901*) and *CirA* (*Xoo3793*) were upregulated and *IroN* genes (*Xoo0394* and *Xoo1784*) were downregulated in both the transcriptome and proteome data. In the proteome data, *FecA* (*Xoo0901*) and *CirA* (*Xoo3793*) upregulation peaked at 30 min (Figure 5A). *IroN* genes that were downregulated at the transcript level were also downregulated at the protein level (Figure 5B). The expression levels of other paralogs of *FecA* and *CirA* genes were comparable with those in the control in the transcriptome as well as the proteome data (Figure S5), indicating that these genes could respond to different pathogenic signals—which were missing in the *in vitro* system—or be pseudogenes.

**Figure 5.**
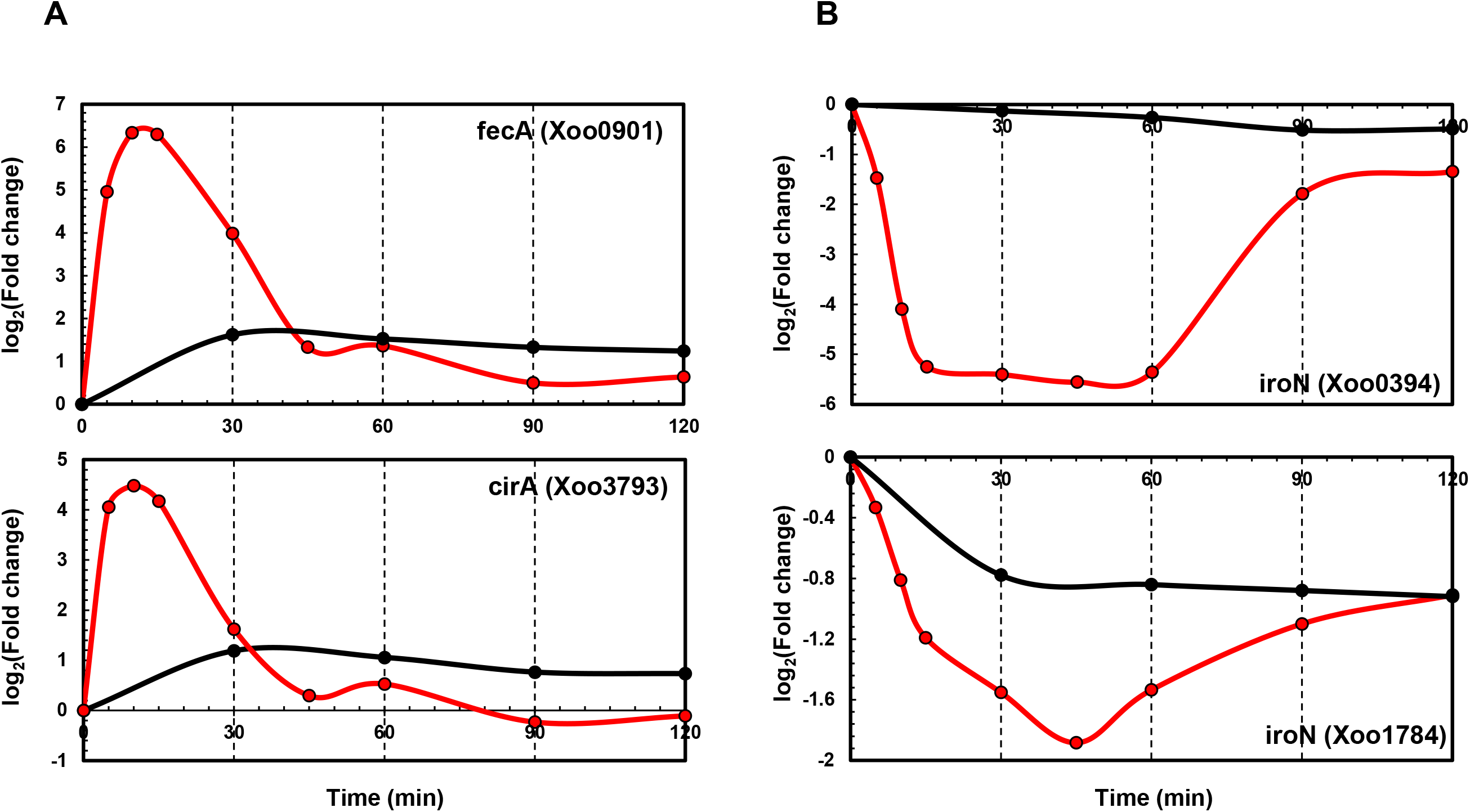
Time-resolved mRNA and protein levels of iron transport-related genes. (A) Time-resolved mRNA (red) and protein (black) expression levels of *FecA* and *CirA* genes. (B) Time-resolved mRNA (red) and protein (black) expression level of *IroN* genes. The Y-axis represents log2(fold-change).

Phosphate uptake genes, i.e., *OprO, PhoX, PstSCAB*, and *PhoU*, were upregulated in the transcriptome (up to 16-fold) at 5-10 min. The proteome data of OprO, PhoX, PstSCAB, and PhoU revealed upregulation up to 3-fold at 30 min (Figure S6).

#### Expression and secretion of effectors

In *Xanthomonas oryzae* pv. *oryzicola*, AvrBs2 suppresses host immunity and promotes disease development (Ullah et al., 1998). In *Xanthomonas campestris* pv. *vesicatoria*, the role of AvrBs3 is that of a transcription activator-like (TAL) effector that activates the expression of plant immunity genes (Kay, Hahn, Marois, Wieduwild, & Bonas, 2009). The secretion of effectors XoAvrBs2 (*Xoo0168*)—a Xoo ortholog of AvrBs2—and XoAvrBs3 (*Xoo2276*)—a Xoo ortholog of AvrBs3—through T3SS was confirmed upon interaction with RLX (S. H. Kim et al., 2011; S. Kim et al., 2013). In addition to the transcriptome and proteome, the expression and secretion of effectors were assessed using dot blots of Xoo cells transformed with the TAP-tagged effector genes (Figure 6).

**Figure 6.**
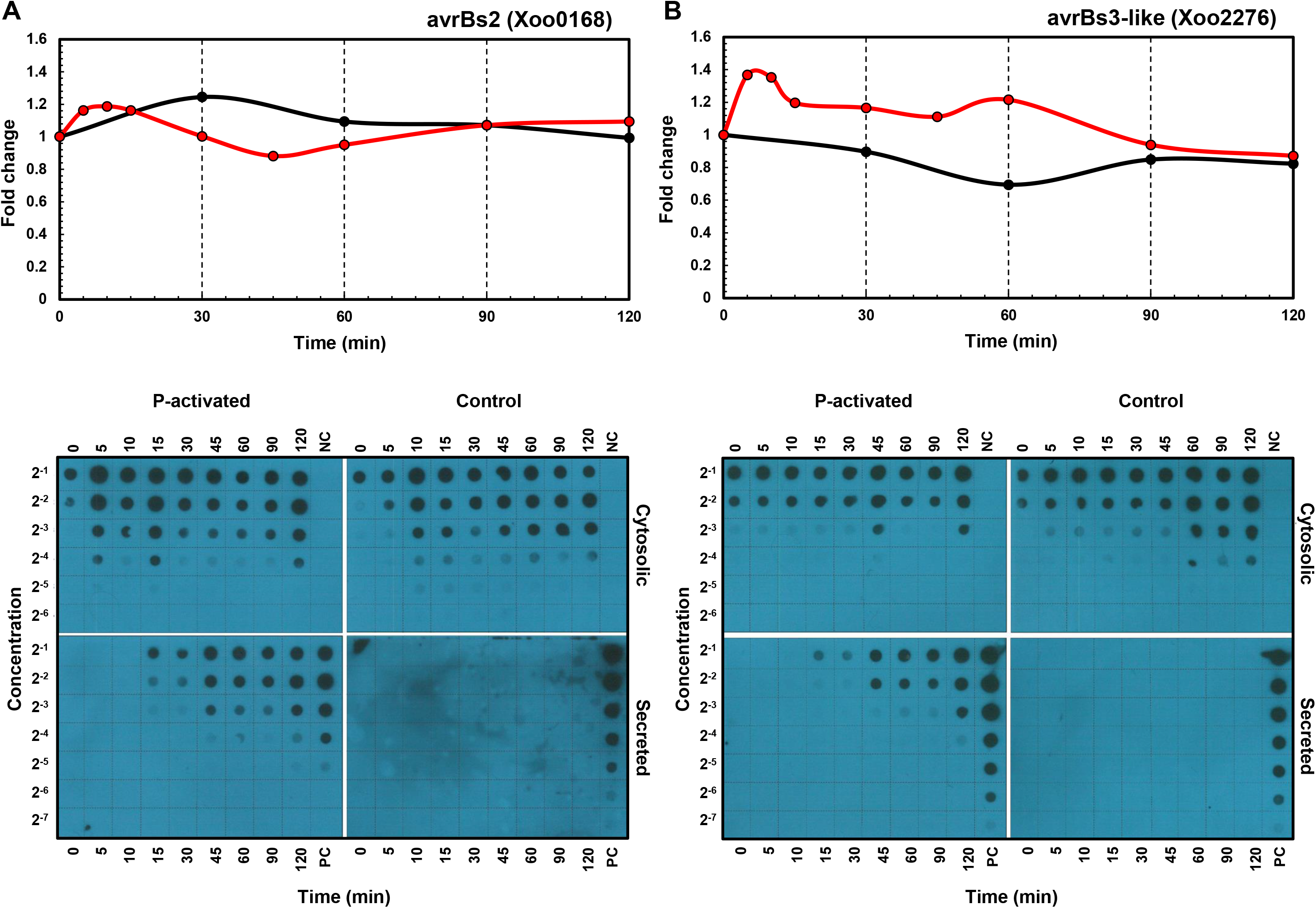
Time-resolved mRNA and protein expression and secretion of effector proteins. The mRNA (red) and protein (black) expression levels of *XoAvrBs2* (A) and *XoAvrBs3* (B) genes in the transcriptome and proteome data (above) and the cytosolic and secreted XoAvrBs2 (A) and XoAvrBs3 (B) proteins in the dot blot data (below). NC, negative control; PC, positive control

The expression of *XoAvrBs2* transcript was upregulated from 5-30 min, while that of the XoAvrBs2 protein was upregulated at 30 min, as is evident from the transcriptome and proteome data. In the dot blots, the levels of TAP-tagged XoAvrBs2 were upregulated by 4-fold at 5 min and the secretion of XoAvrBs2 was detected as early as 15 min. The expression of *XoAvrBs3* transcript was upregulated and peaked from 5-30 min, whereas that of XoAvrBs3 protein was maintained at almost the same—or slightly lower levels—at 60 min in the transcriptome and proteome data. Dot blots revealed the secretion of XoAvrBs3 from 15 min.

## Discussion

Plant pathogens could exhibit complex responses to the initial interactions with the host under varying biotic and abiotic conditions at the site of infection. The varied immediate response of the plant pathogen is important for successful infection. In this study, we analyzed the immediate time-resolved response of Xoo cells from the initial interaction with rice in terms of gene expression at both the mRNA and protein levels using an *in vitro* assay system, wherein RLX mimicked the damaged rice leaf tissue (Scheme 1).

Of all the predicted open reading frames in Xoo, we quantified approximately 83% and 47% of the mRNAs and proteins, respectively, in a time-dependent manner. A good correspondence of overall expression pattern was observed between the mRNA and protein levels of genes. The expression pattern was more synchronized for genes closely located in the genome (Figure 4 and S4); this could be attributed to the polycistronic gene structure in bacteria. With respect to the rapidity of pathogenic gene expression in response to the signals triggered upon interaction with the host, both transcriptional and translational machineries responded immediately to the interaction, at the earliest assessed time points of 5 and 30 min in the transcriptome and proteome data. The secretion of effectors was also found at 15 min in dot blots. Both mRNAs and proteins presented the greatest variation in their levels during the initial 30 min.

The Xoo genes exhibiting the most rapid responses to the initial interaction with RLX, included genes associated with cell motility and inorganic ion uptake, and genes coding for effector molecules. All these three functional categories of genes are closely related to the early stages of pathogenesis. The genes associated with cell motility could be responsible for the migration and accumulation of the Xoo cells at the site of infection via the damaged xylem tissues or exposed hydathodes in rice. As inorganic ions, such as iron and phosphate ions, function as essential cofactors in all living organisms, bacterial pathogens and host rice cells compete to obtain and secure the limited resources available. Effector molecules are more directly related to pathogenesis and are injected by the Xoo into the rice cells via pili-like T3SS; these molecules modulate the immune system of rice.

The earliest available time-point for comparing the gene expressions in terms of mRNA and protein levels was 30 min after the RLX treatment. At 30 min, 290 mRNAs were upregulated by more than 2-fold in the transcriptome, compared with 93 proteins in the proteome (Table S4), and the average fold change in the expression of the upregulated genes in the transcriptome and proteome was similar, i.e., 3.3-fold and 3.4-fold, respectively (Table S6). The number of upregulated transcripts is much higher than—even though we figure into our calculations the higher coverage of quantified genes in transcriptome than in proteome—that of the proteins at 30 min, which indicates that not all the upregulated mRNAs are simultaneously translated to proteins.

The difference in the gene expression in terms of the mRNA and protein levels at a given time point suggests the existence of an additional translational regulation step—after the transcriptional regulation—although transcription and translation can occur simultaneously in bacteria. For some genes, the mRNA and protein expression peaks were observed at varying time points. Bacterial pathogenesis appears to involve a fine-tuning mechanism after transcription, which could help adjust the expression of the early-responsive genes under varying biotic and abiotic environmental conditions.

Genes related to flagella and cell motility are clustered in the bacterial genome. In Xoo, three clusters are observed, i.e., I-A, I-B, and II. On superimposing the mRNA and protein levels in time, the expression of mRNAs and proteins peaked at 30 mins for genes in the clusters I-A and I-B, the expression of mRNAs and proteins peaked at 30 and 90 min, respectively, for genes in cluster II. A translational regulation step might be involved that determines the time for the translation of specific mRNAs—coded by flagella and cell motility-related genes—into proteins depending on the priority of each gene.

The hierarchy of the expression of flagella and cell motility-related genes has been extensively studied in the transcriptome of *E. coli*, where sigma and anti-sigma factors are known to be the key transcriptional regulators (Mukherjee & Kearns, 2014). The overall organization of flagella and chemotaxis-related genes is different between *E. coli* and Xoo (Figure S7). In Xoo, the cluster of *fliE-R* genes (cluster I-B) is positioned just downstream to the *flgB-L* genes (cluster I-A), whereas in *E. coli*, chemotaxis genes are present between the two. *flhDC* genes do not have any orthologs in Xoo, and a different transcriptional regulator, i.e., the *fleQ* gene, is present. Based on the nomenclature used in *E. coli*, several Xoo genes classified as Class III genes are located at different positions and their expression regulation is also different. In Xoo, ribosomes or other translation factors might recognize certain unknown priority signals in mRNA transcripts for the translational regulation.

The immediate upregulation of inorganic ion uptake genes, especially iron uptake genes, may aid Xoo cells to obtain the essential cofactor ions, when they are competition with the host cells (Liu, Kong, Wu, & Ling, 2020). The iron ion is essential for photosynthesis and respiration and needed by many redox enzymes for most organisms on Earth. Pathogens commonly use iron chelating molecules or siderophores to derive this scarce inorganic cofactor from the hosts. The leakage of iron ions from damaged leaf tissues might serve as an important signal of initiating infection and might provide an opportunity to secure essential iron ions for the Xoo cells.

In comparison with the flagella and cell motility-related genes that are closely clustered in the genome and exhibit coordinated expression levels, the ion uptake genes are dispersed across the genome and are independently expressed (Figure S8). The separation of these genes facilitates independent regulation. Compared with the flagella and motility-related genes, the inorganic ion uptake genes exhibited great variation in the mRNA levels but similar protein expression levels (Figure 3).

Effectors are key molecules for pathogenicity that modulate the host immune responses after infection. The protein expression and secretion of the Xoo effectors XoAvrBs2 and XoAvrBs3 were assessed using dot blots, which enabled the monitoring of effector proteins at the early stages of the initial interaction, i.e., within 30 min of RLX treatment. The secretion of effectors was observed from 15 min after application of the pathogenic stimulus. In the P-activated proteome data, the expression of effector proteins was maintained at a similar level to that at 0 min; this may be attributed to the similar protein synthesis and secretion rates.

The effector genes are also dispersed across the genome, like the ion uptake genes (Figure S9). Interestingly, transposase genes are located close to the effector genes; these may aid the effector genes to transpose through the bacterial genome and plasmids. The expression of XoAvrBs2 was upregulated 5 min after RLX treatment, and its secretion was detected in the culture medium at 15 min. Although the cellular protein level was upregulated by approximately 20% in the proteome at 30 min, the dot blot showed an upregulation of 4-fold at 5 min. The secreted XoAvrBs2 exhibited a 16-fold increase at 120 min. In case of XoAvrBs3, the cellular protein level decreased by approximately 20% in the proteome at 60 min, whereas the dot blot revealed a 16-fold increase in XoAvrBs3 secretion at 120 min. The experimental methods for proteome and dot blots analysis are different (S. H. Kim et al., 2011; S. Kim et al., 2013). For the dot blots analysis, TAP-tagged *XoAvrBs2* and *XoAvrBs3* genes were introduced into Xoo cells via the plasmid having the endogenous promotor and expressed from the plasmid. In the proteome analysis, XoAvrBs2 and XoAvrBs3 proteins were expressed from the endogenous *XoAvrBs2* and *XoAvrBs3* genes in the Xoo genome.

Gene expression involves sequential transcription and translation. In bacteria, with respect to post-transcriptional and post-translational modification, mRNAs without a cap at 5’ end and a poly A tail at 3’ end have a short half-life—as short as few minutes—and proteins undergo only limited post-translational modifications. The limited post-translational modifications in bacterial proteins impose a pressure on a nascent protein from the ribosome to take a functional form immediately. The present study revealed that genes related to cell motility and inorganic ion uptake, and genes coding for effector molecules of Xoo are the first to respond to the initial interactions with RLX, and play an essential role in the following processes (Figure 7): (1) invasion of the Xoo cells into the rice leaf tissues, (2) securing the limited cofactors, and (3) modulating the immune responses of the host to favor pathogenesis. This combined analysis of the time-resolved transcriptome and proteome of Xoo during the initial interaction with rice tissues provides valuable insights into the pathogenic mechanism of Xoo.

**Figure 7.**
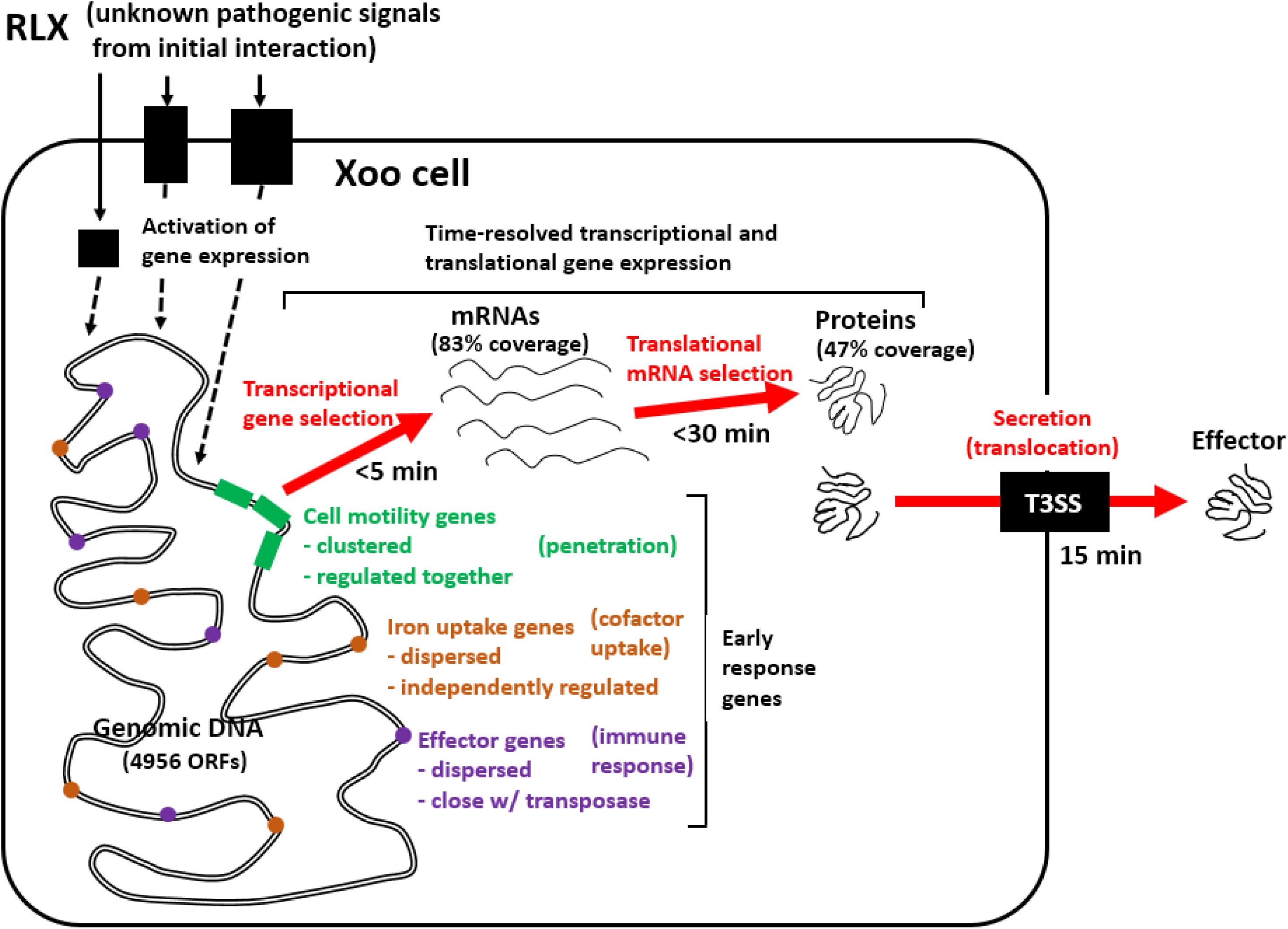
Schematic representation of genome-wide pathogenic gene expression and effector secretion via in vitro assay system. Early response Xoo genes from the initial interaction with RLX include genes related to cell motility, iron uptake, and effector, of which expression are upregulated as early as 5 min in mRNAs and 30 min in proteins and effector secretion is found since 15 min.

## Materials and Methods

### Bacterial strain and culture conditions

*Xanthomonas oryzae* pv. *oryzae* (Xoo) strain KACC10331, consisting of 4,941,439 nucleotides and 4,733 open reading frames, without any apparent autonomous plasmids, was obtained from the Korean Agricultural Collection Center (KACC) (Lee et al., 2005). The bacteria were cultured in nutrient broth (Difco, Detroit, MI, USA) or Yeast Glucose Cm Agar (YGC) (1% yeast extract, 2% D-(+)-glucose, 2% CaCO3, and 1.5% agar) at 28°C.

### Construction of expression vector and transformation of Xoo

The effector genes *XoAvrBs2* (*Xoo0168*) and *XoAvrBs3*—including the promoter region (from −149 and −750 bp to the start codon of the respective gene)—were amplified by PCR and ligated into the pGEM®-T Easy Vector (Promega). The cloned sequences were verified and then digested with *KpnI* and *SacI*, and the products were ligated into the pHM1-XTAP-Tgap vector. The recombinant vectors were purified and introduced into Xoo strain KACC10331 by electroporation, using Gene Pulser II (Bio-Rad, Hercules, CA) with a 0.2 cm-gap cuvette at 2.5 kV cm^-1^. Xoo cells were then diluted immediately with 1 mL Super Optimal Broth (SOC) medium and incubated at 28°C with agitation for 2 h. Cells were then recovered from the culture medium and plated on nutrient broth agar plates containing 50 μL mL^-1^ spectinomycin and incubated at 28°C for 4 d.

The transformants were cultured in 100 mL of nutrient broth up to the mid-exponential phase (A_600_ = 0.5). Cells were harvested by centrifuging 1 mL of the cell culture at 12,000 rpm and 4°C for 5 min. Harvested cells were washed once with phosphate-buffered saline (PBS) at pH 7.2, resuspended in 200 μL PBS, and then sonicated. Protein samples were serially diluted using 2 M urea in PBS in 96-well plates and then transferred to a polyvinylidene difluoride membrane (PVDF; 0.2 μm, Bio-Rad) using a 96-well vacuum dot-blotter (Bio-Rad). The membrane was then washed thrice with PBS, blocked with 5% skim milk for 30 min, and subjected to a one-step immuno-affinity reaction using the rabbit peroxidase-anti-peroxidase soluble complex antibody (Sigma-Aldrich, St. Louis, USA). The membrane was developed, and bound antibodies were detected by chemiluminescence.

### Treatment of rice leaf extract for proteome analysis

*Oryza sativa* L. cv. Milyang 23, a Xoo-susceptible rice cultivar, was used for performing proteome analysis. Rice plants were grown in a paddy field at Jeonju in South Korea (35°49’52.0”N 127°03’55.6”E) until panicle initiation (approximately 8 to 9 weeks). Forty clumped rice leaves were harvested and homogenized with liquid nitrogen using a mortar and pestle. One-gram aliquots of the resulting homogenate (RLX) were transferred to Eppendorf tubes and stored at −80°C. Xoo was cultured (100 mL) in nutrient broth up to the mid-exponential phase (A600 = 0.5) in a shaking incubator at 28°C and 200 rpm, and RLX (2 g) was then added to the culture medium. The culture (100 mL) was subjected to sequential filtration through a gauze, 40-μm nylon cell strainer (FALON, New York, USA), and 5-μm syringe filter (Sartorius, Germany) to remove RLX (0, 30, 60, 90, and 120 min after RLX addition). The filtered culture (100 mL) was centrifuged at 10,000 ×*g* and 4°C for 10 min. Duplicate samples were obtained for each time point from two independent experiments.

### Sample preparation for proteome analysis

The harvested samples were lysed in a lysis buffer (9 M urea prepared in 20 mM HEPES (pH 7.5), supplemented with protease inhibitor cocktail (Complete mini, Roche) and phosphatase inhibitor (PhosSTOP, Sigma-Aldrich)) and sonicated on ice. The exact amount of proteins in each sample was determined using the bicinchoninic acid assay. Protein integrity was confirmed by SDS-PAGE and 200 μg of protein from each sample was used for analysis. The disulfide bonds were reduced by treatment with 10 mM dithiothreitol for 1 h, and incubation with 30 mM iodoacetamide (30 min in the dark) was performed to alkylate free sulfhydryl functional groups. Samples were diluted with triethylammonium bicarbonate buffer (pH 8.0) in a manner such that the final urea concentration was 1.5 M. Proteins were digested using MS grade trypsin (Thermo Fisher Scientific) at a protein to enzyme ratio of 50:1 for 12 h at 37°C. The reaction was quenched by lowering the sample pH (<3) using trifluoroacetic acid. The obtained peptides were desalted using a C18 spin column (Harvard) to remove salts and other contaminants, and the purified peptides were dried. Then, they were isotopically labeled using the 10-plex tandem mass tag (TMT, Thermo Fisher Scientific), as per the manufacturer’s protocol. The reaction was allowed to continue for 2 h at room temperature and TMT-labeled samples were subsequently dried in a SpeedVac concentrator. Chemical labeling with TMT was confirmed by liquid chromatography-tandem mass spectrometry (LC-MS/MS), and the samples were pooled and fractionated using a basic reverse phase liquid chromatography (RPLC) system. The pooled TMT-labeled peptide mixture was resuspended in 10 mM ammonium formate and fractionated into 12 fractions using a C18 column (C18, 5 μm pore size, 4.6 mm × 250 mm, XBridge, Waters). The fractionated peptides were dried and stored at −80°C until LC-MS/MS analysis.

### LC-MS/MS and database search

Each fractionated peptide sample was analyzed using an Orbitrap Fusion™ Lumos™ Tribrid™ Mass Spectrometer coupled with the Easy-nLC 1200 nano-flow liquid chromatography system (Thermo Fisher Scientific). The dried peptides were reconstituted using 0.1% formic acid and loaded on a C18 trap column. Peptides were resolved using a linear gradient solvent B (0.1% formic acid in 95% acetonitrile) and analyzed by high resolution mass spectrometry in the data-dependent acquisition mode. MS1 and MS2 were acquired for the precursor and the peptide fragmentation ions, respectively. MS1 scans were measured at a resolution of 120,000 and an *m/z* of 200. MS2 scans were acquired following the fragmentation of precursor ions by high-energy collisional dissociation (HCD) and were detected at a mass resolution of 50,000 and an *m/z* of 200. Dynamic exclusion was used to reduce redundant fragmentation of the same ions. The obtained mass spectrometry data were analyzed using the MaxQuant software (Tyanova, Temu, & Cox, 2016). Raw MS data were searched against the Xoo proteome in UniProt database. Carbamidomethylation of cysteine and 10-plex TMT modification of lysine and N-terminals were set as static modifications, whereas oxidation of methionine was set as a variable modification. False discovery rates at the levels of protein and peptide-spectrum matches were set at 0.01. The raw MS data and MaxQuant search results have been submitted to ProteomeXchange (project accession: PXD020135, reviewer access with username: and password: mzn76I1O).

### Proteomic data analysis

The contaminant and reverse identified proteins were removed from the MaxQuant data. Proteins identified in both replicates were pooled for quantile normalization. The normalized values for the replicates were subjected to supervised hierarchical clustering and principal component analysis, using Perseus (Tyanova, Temu, Sinitcyn, et al., 2016), and the results were depicted in the form of a multi-scatter plot.

### STRING map analysis

The list of proteins in the selected patterns from the hierarchical clusters was uploaded on the STRING database (https://string-db.org/) to analyze the protein interaction maps. The clusters of proteins in the blue and red classes, which show the similar time-resolved expression patterns, were analyzed, and the list including the gene names with the selected organism was inputted in the multiple proteins search setting. The number of nodes and edges were automatically calculated based on Xoo genes with a PPI enrichment p-value of 20.9E-10. Figures were downloaded in the PNG file format for visualization.

### RNA-Seq and data analysis

In addition to the previously obtained RNA-Seq data for P-activated Xoo cells, RNA-Seq data at 90 and 120 min were obtained to correspond with the proteome data at these time points. RNA-Seq and data analysis were performed as previously described (S. Kim et al., 2016). Briefly, total RNA from samples was used to generate sequencing libraries, from which ribosomal RNA was removed using MICROBExpress Bacterial mRNA Enrichment Kit (Ambion, Austin, TX, USA), and enriched mRNA was prepared using Illumina TruSeq RNA Sample Preparation Kit (Illumina, San Diego, CA, USA). The RNA obtained after fragmentation was used to generate cDNA fragments, which were sequenced using Illumina Genome Analyzer IIx and mapped to the reference genome sequence (http://www.ncbi.nlm.nih.gov/nuccore/58579623?report=fasta) using CLC Genomics Workbench 4.0 (CLC bio, Aarhus, Denmark). Relative transcript abundance was calculated based on the number of reads per kilobase per million mapped sequence reads (RPKM).

### Analysis of time-resolved continuous mRNA and protein expression

The RPKM values of the mRNAs in the transcriptome and TMT intensities of the proteins in the proteome corresponded to the observed expression level of each gene at a given time point. To facilitate the comparison of gene expression levels, the observed expression level at each time point was converted to fold change in gene expression, by dividing the expression level at a given time point by the initial expression level (0 min) of the same gene. The fold change in the time-resolved expression levels of a given gene during the two hours following the RLX treatment was fitted to a curve and analyzed using non-linear regression by GraphPad Prism (version 3.02 for Windows, GraphPad Software, San Diego California USA, www.graphpad.com), and the continuous time-dependent changes in the mRNA and protein expression levels were determined using the fitted curve.

### Analysis of the correspondence between mRNA and protein expression levels

For the accurate comparison of the mRNA and protein expression levels in P-activated Xoo cells, the expression levels were corrected by comparing with those in the control cells at each time point. The fold change in mRNA and protein expression in P-activated Xoo cells at a given time point was divided by that of the control cells at the same time point. For analyzing the correspondence between the mRNA and protein levels of each gene in the transcriptome and proteome, the resulting control-corrected fold change values of mRNAs were compared with those of the proteins for a given time point as the gene expression level of mRNAs and proteins.

## Declarations

### Available data and material

The raw MS data and MaxQuant search results are available at ProteomeXchange (Project accession: PXD020135, Reviewer access with username: and password: mzn76I1O)

### Author Contributions

Investigation, Seunghwan Kim, Wooyoung Eric Jang, Min-Sik Kim, Jeong-Gu Kim, and Lin-Woo Kang; writing, Seunghwan Kim, Wooyoung Eric Jang, Min-Sik Kim, Jeong-Gu Kim, and Lin-Woo Kang; methodology, Seunghwan Kim, Wooyoung Eric Jang, Min-Sik Kim, Jeong-Gu Kim, and Lin-Woo Kang; funding acquisition, Jeong-Gu Kim, and Lin-Woo Kang; supervision, Min-Sik Kim, Jeong-Gu Kim, and Lin-Woo Kang. All authors have read and agreed to the published version of the manuscript.

## Acknowledgments

This study was supported by the Cooperative Research Program for Agriculture Science & Technology Development (Project No. PJ01327002020), Rural Development Administration and by the 2021 RDA Fellowship Program of National Institute of Agricultural Sciences, Rural Development Administration, Republic of Korea.

## Abbreviations

Xoo: *Xanthomonas oryzae* pv. *oryzae*
RLX: rice leaf homogenate
KACC: Korean Agricultural Collection Center
P-activated: Pathogenicity-activated
LC-MS/MS: liquid chromatography-tandem mass spectrometry
COG: Clusters of Orthologous Groups
RPKM: Reads per kilobase per million mapped reads

