## Supplementary_Figs_and_legends for "Combined analysis of the time-resolved transcriptome and proteome of plant pathogen *Xanthomonas oryzae* pv. *oryzae*"

### Supplementary data

#### Supplementary figure legends

**Figure S1. Detailed analysis of the proteome data of pathogenicity-activated Xoo cells** (A) Number of proteins identified from the high-resolution mass spectrometry-based quantitative proteomic analysis and (B) coverage percentage of the protein sequence of the identified proteins (median sequence coverage was ~24%). (C) Number of identified peptides and peptide-spectrum matches. (D) Quantile normalization box plot. (E) Multi-scatter plot for analyzing the correlations between samples.

**Figure S2. Comparative analysis of the proteome of the samples** (A) Quantile normalization box plot. The quantitative proteomic values were normalized using quantile normalization. (B) Multi-scatter plot for analyzing the correlation between samples. Pairwise comparisons of protein expression levels in all samples are presented as a multi-scatter plot. Pearson's correlation coefficients of 0.98-0.99 were obtained. (C) Principal component analysis of Xoo proteins revealed close relationships among the proteomes of all controls and changes in proteome concentrations upon RLX treatment.

**Figure S3. Analysis and comparison of gene expression patterns in the datasets.** The STRING maps (Benjamin-Hochberg at FDR 0.05) of selected cluster profile patterns with (A) downregulated (blue box) and (B) upregulated (red box) genes are shown.

**Figure S4. Comparison of time-resolved mRNA and protein levels of cell motility-related genes.** (A) Time-resolved mRNA levels of flagellar biosynthesis and chemotaxis-related genes for pathogenicity-activated Xoo cells. All genes in groups I-A, I-B, and II exhibited the lowest expression level at 5 min and the highest expression level at 30 min. (B) Time-resolved protein levels of flagellar biosynthesis and chemotaxis-related genes in pathogenicity-activated Xoo cells. Genes in groups I-A and I-B exhibited the highest expression level at 30 min, whereas those in group II exhibited the highest expression at 90 min.

**Figure S5. Time-resolved mRNA and protein expression of iron uptake-related genes.** Time-resolved mRNA (red) and protein (black) expression levels of iron uptake-related genes (A) *FecA* and (B) *CirA*. The Y-axis represents $\log_2$(fold change).

**Figure S6. Gene cluster and time-resolved mRNA and protein expression levels of phosphate uptake-related genes.** (A) Gene cluster of the phosphate uptake regulation genes (*OprO-PhoX-PstSCAB-PhoU*). (B) Time-resolved mRNA and protein expression levels of phosphate uptake-related genes *OprO*, *PhoX*, *PstSCAB*, and *PhoU.* The line colors correspond to those of the arrows in (A).

**Figure S7. Gene clusters of cell motility-related genes in *E. coli* and Xoo and time-resolved mRNA and protein expressions of Xoo genes.** Gene cluster of cell motility-related genes of (A) *E. coli* and (B) Xoo. (C) Time-resolved mRNA and protein expression of cell motility-related genes in Xoo. The cell motility-related genes in *E. coli* are classified as class I (red), II (yellow), III (blue), and II+III (green). The ortholog genes in Xoo are indicated using the same color. For the proteome data, * indicates the expression peak at 30 min; **, between 30 and 90 min; ***, at 90 min. The Y-axis represents $\log_2$(fold change).

**Figure S8. Gene clusters and time-resolved mRNA and protein expressions of iron uptake-related genes in Xoo.** (A) Gene cluster of iron uptake-related genes, labelled in blue. (B) Time-resolved mRNA and protein expression levels of iron uptake genes. Low to high expression is indicated by a change in color from blue to red. Black cells indicate undetected expression.

**Figure S9. Gene clusters and time-resolved mRNA and protein expressions of genes coding for effector molecules.** (A) Gene cluster of effector genes, indicated in red. (B) Time-resolved mRNA and protein expressions of effector genes. Genes with available time-resolved proteomic data are labelled with *. Of the *avrXa7* and *avrXa3* genes, the mRNA levels of only the former were measured, indicated by +. Transposase genes are indicated in blue.

#### Supplementary Table legends

**Table S1. The raw data of time-resolved expression levels of mRNAs and proteins in the transcriptome and proteome of P-activated and control Xoo cells.** The RPKM values of mRNAs and TMT intensities of proteins were obtained from independent duplicated transcriptome and proteome analyses, respectively, representing the expression levels of mRNAs and proteins of each gene. The gene products that were not identified or measured are indicated by blank cells.

**Table S2. The list of selected genes with similar protein expression patterns in the time-supervised hierarchical clustering.** The list of genes marked with blue (downregulated) and red (upregulated) rectangles in Figure 1, which presented distinct synchronized protein expression patterns. Of the 40 flagellum-related KEGG annotated genes, 21 were included in the list, of which 20 genes were upregulated and one gene was downregulated in the proteome.

**Table S3. The fold change of time-resolved expression levels of mRNAs and proteins in the transcriptome and proteome of P-activated and control Xoo cells.** The fold change (compared to 0 time) of expression levels is calculated by dividing the expression levels of mRNAs and proteins at a given time point by that at 0 min. Cells are colored based on the expression level; low to high expression is indicated by a change in color from blue to red, with white indicating a fold change of 1.

**Table S4. The control-corrected fold change of time-resolved expression levels of mRNAs and proteins in the transcriptome and proteome.** For the convenient comparison of the mRNA and protein expression levels, the expression levels of mRNAs and proteins in P-activated Xoo cells at each time point was divided by that of the control cells (RLX-untreated Xoo cells) at the same time point. The number and percentage of genes exhibiting more than 20%, 50%, and 200% (2-fold) up- and downregulation are indicated at the bottom of the table.

**Table S5. The number of consistently up- and downregulated genes for 120 min.** The numbers of more than 20%, 50%, and 2-fold up- and downregulated genes are measured in Xoo cells for the initial 20 min since the first interactions with rice.

**Table S6. The list of genes having 2-fold upregulated proteins and mRNAs at 30 min.** The lists contain genes having more than 2-fold upregulated proteins at 30 min in both duplicate proteome datasets (S6-1), genes having more than 2-fold upregulated proteins at 30 min in the proteome repetition dataset 1 (S6-2) and 2 (S6-3), and genes having more than 2-fold upregulated mRNAs at 30 min in the transcriptome dataset (S6-4).

**Table S7. The list of genes having 50% downregulated proteins at 30 min in the duplicate proteome datasets.** The list contains genes having more than 50% downregulated proteins at 30 min in both proteome datasets.

### Supplementary data

### Figure S1

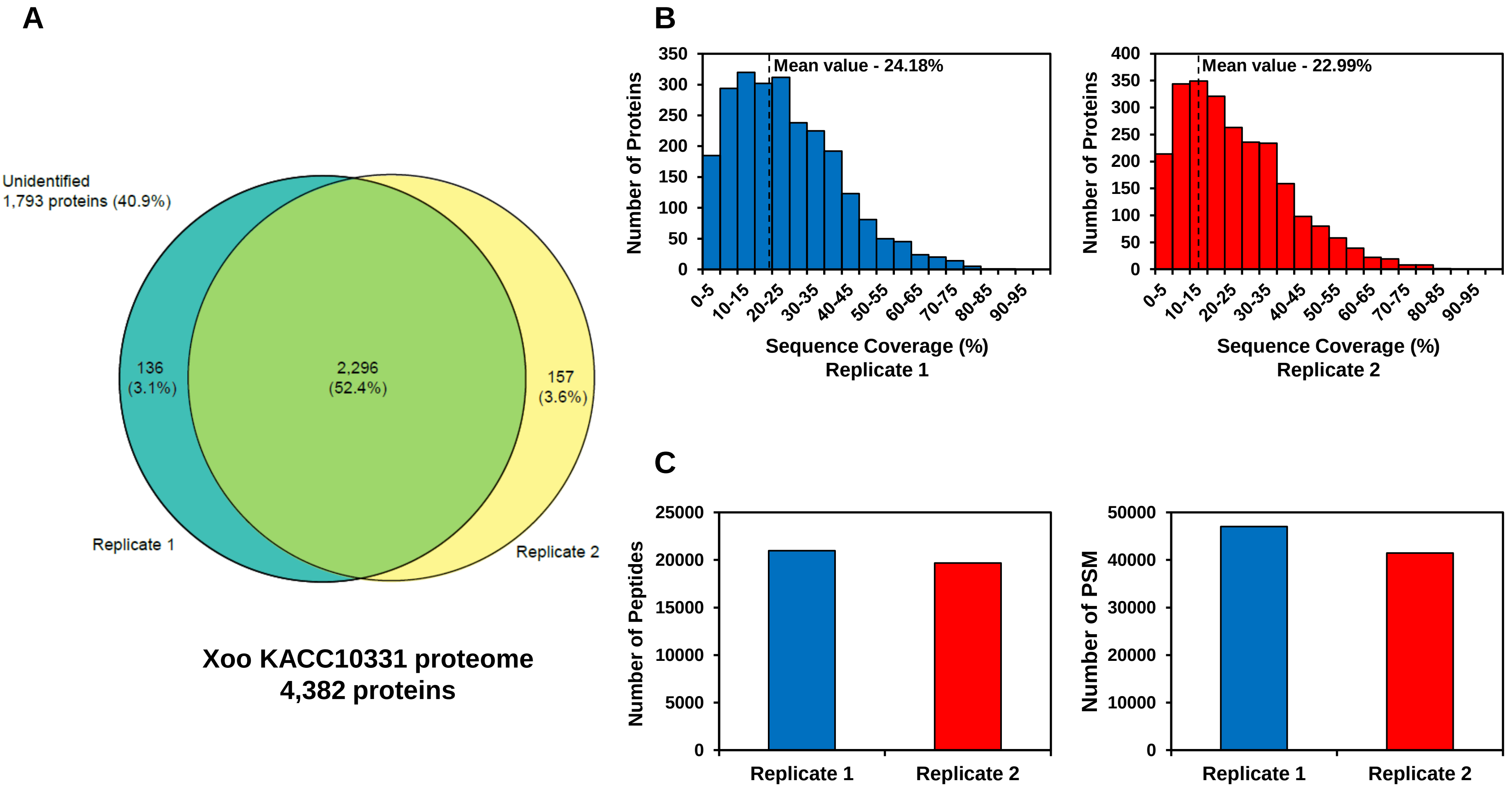

A
Unidentified
1,793 proteins (40.9%)
136
(3.1%)
2,296
(52.4%)
157
(3.6%)
Replicate 1
Replicate 2
Xoo KACC10331 proteome
4,382 proteins
B
Mean value - 24.18%
Number of Proteins
Sequence Coverage (%)
Replicate 1
Mean value - 22.99%
Number of Proteins
Sequence Coverage (%)
Replicate 2
C
Number of Peptides
Replicate 1
Replicate 2
Number of PSM
Replicate 1
Replicate 2

### Figure S2

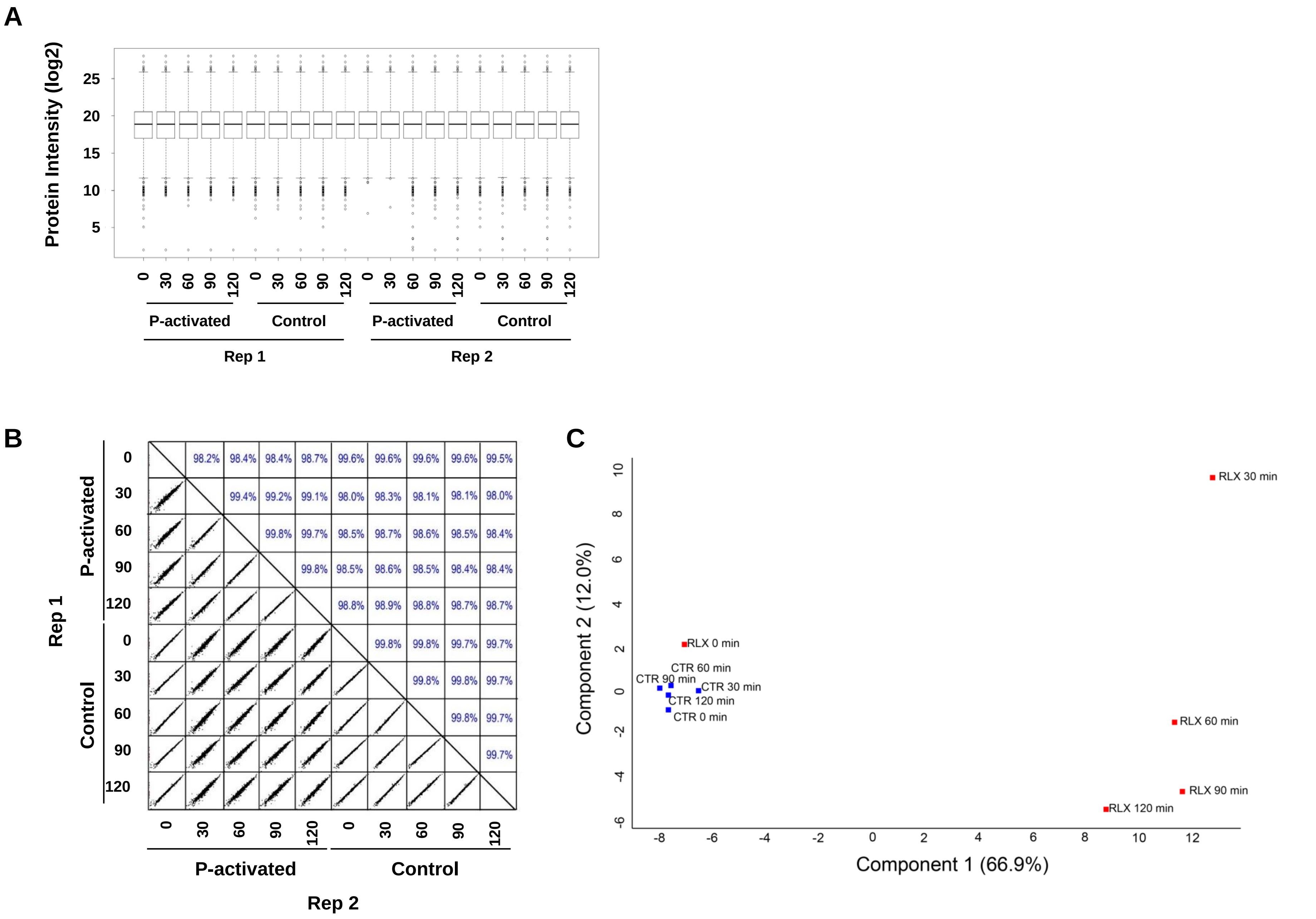
A
Protein Intensity (log2)
25
20
15
10
5
0 30 60 90 120 0 30 60 90 120 0 30 60 90 120 0 30 60 90 120
P-activated
Control
P-activated
Control
Rep 1
Rep 2
B
98.2% 98.4% 98.4% 98.7% 99.6% 99.6% 99.6% 99.6% 99.5%
99.4% 99.2% 99.1% 98.0% 98.3% 98.1% 98.1% 98.0%
99.8% 99.7% 98.5% 98.7% 98.6% 98.5% 98.4%
99.8% 98.5% 98.6% 98.5% 98.4% 98.4%
98.8% 98.9% 98.8% 98.7% 98.7%
99.8% 99.8% 99.7% 99.7%
99.8% 99.8% 99.7%
99.8% 99.7%
99.7%
P-activated
Control
Rep 1
0 30 60 90 120 0 30 60 90 120
P-activated
Control
Rep 2
C
RLX 30 min
RLX 0 min
CTR 60 min
CTR 90 min
CTR 30 min
CTR 120 min
CTR 0 min
RLX 60 min
RLX 90 min
RLX 120 min
Component 2 (12.0%)
10
8
6
4
2
0
-2
-4
-6
-8 -6 -4 -2 0 2 4 6 8 10 12
Component 1 (66.9%)

### Figure S3

**A**

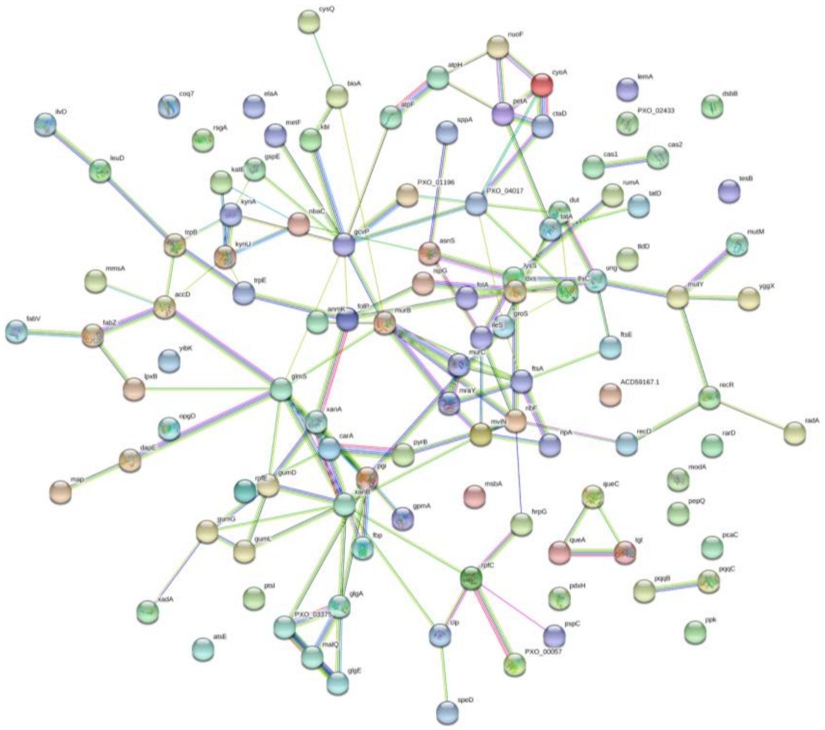

**B**

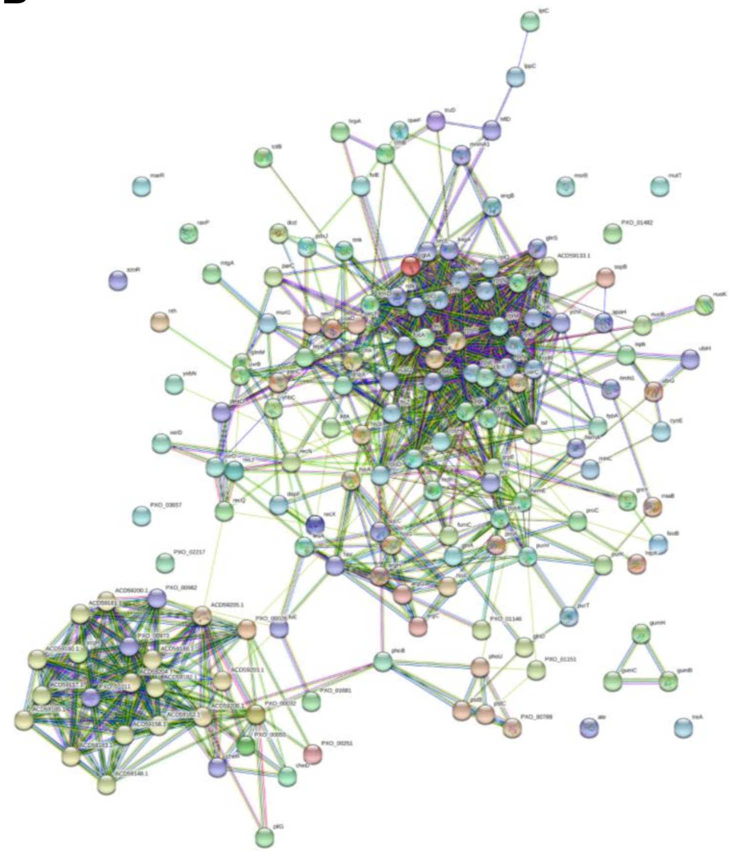

Figure S4

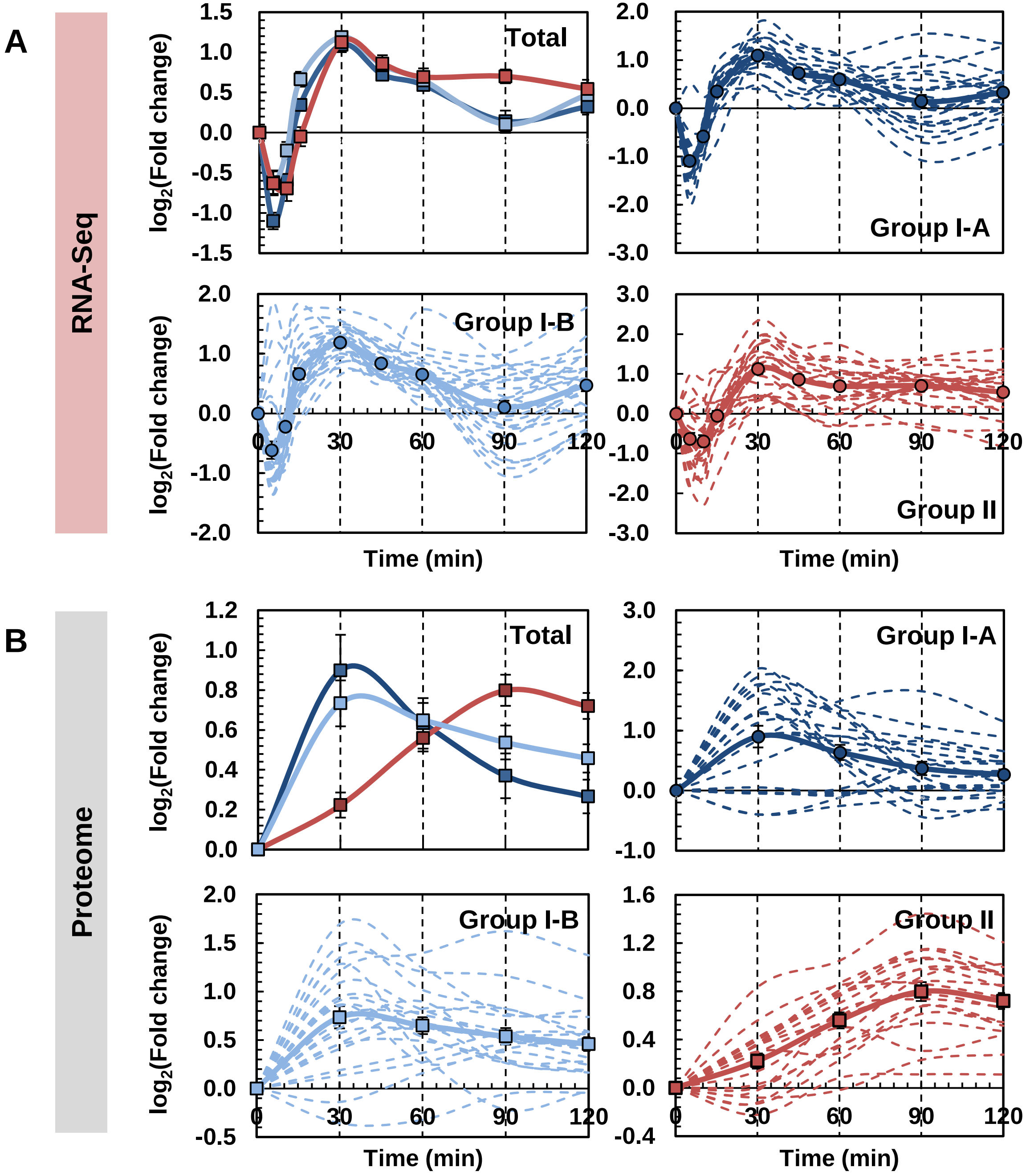
A
RNA-Seq
Total
Group I-A
Group I-B
Group II
log2(Fold change)
Time (min)
B
Proteome
Total
Group I-A
Group I-B
Group II
log2(Fold change)
Time (min)

### Figure S5

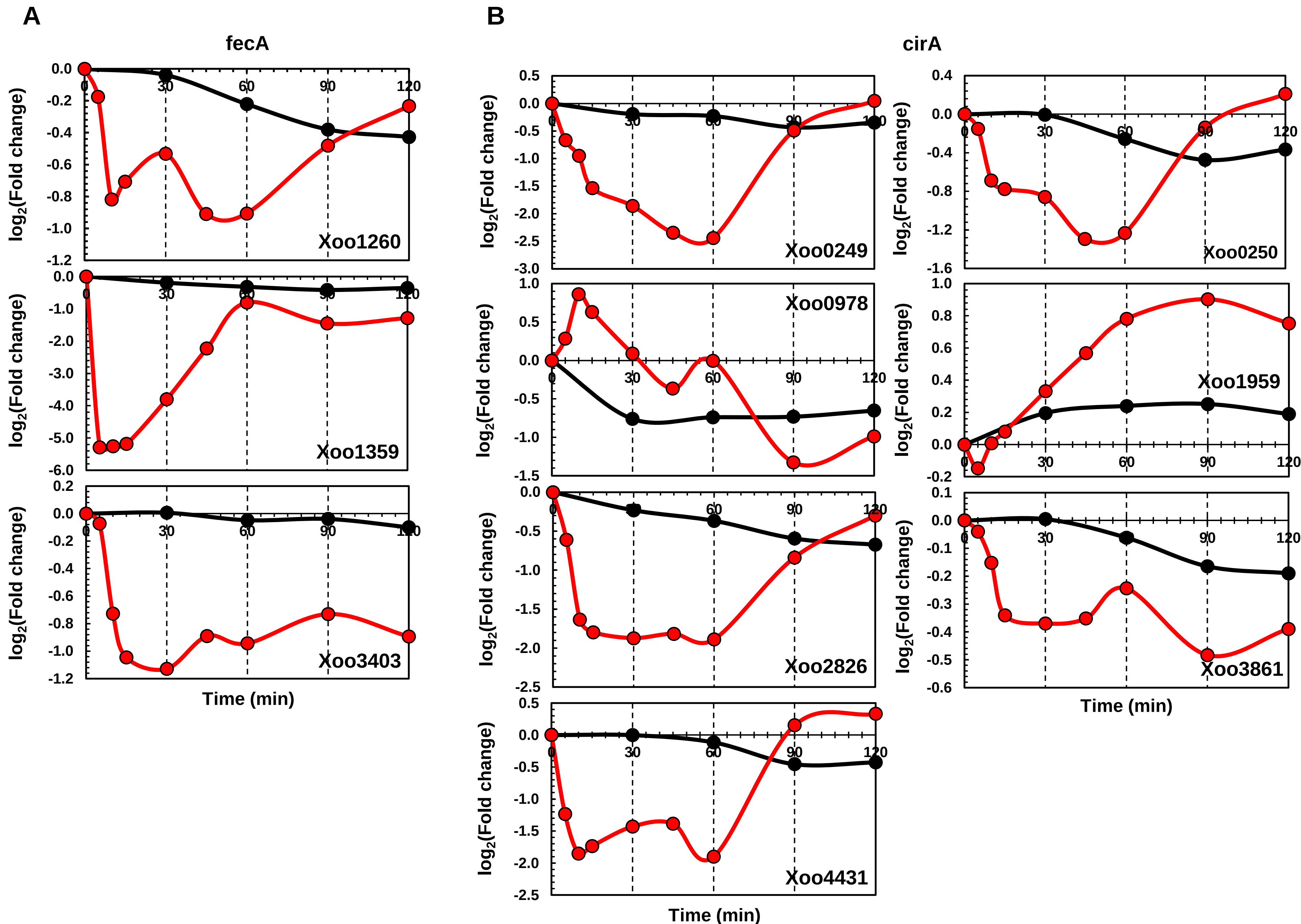

A
B
fecA
cirA
log2(Fold change)
Time (min)
Xoo1260
Xoo1359
Xoo3403
Xoo0249
Xoo0250
Xoo0978
Xoo1959
Xoo2826
Xoo3861
Xoo4431

### Figure S6

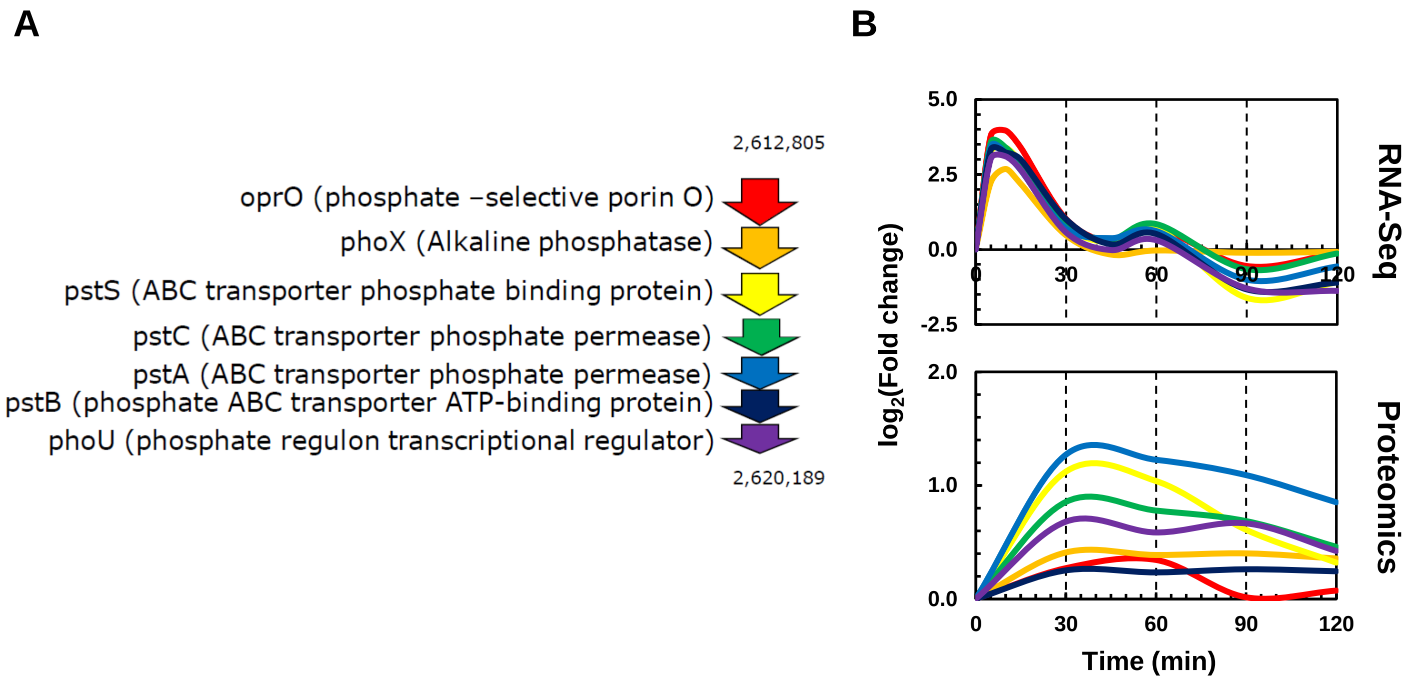
A
B
2,612,805
oprO (phosphate –selective porin O)
phoX (Alkaline phosphatase)
pstS (ABC transporter phosphate binding protein)
pstC (ABC transporter phosphate permease)
pstA (ABC transporter phosphate permease)
pstB (phosphate ABC transporter ATP-binding protein)
phoU (phosphate regulon transcriptional regulator)
2,620,189
5.0
2.5
0.0
-2.5
2.0
1.0
0.0
0
30
60
90
120
log2(Fold change)
RNA-Seq
Proteomics
Time (min)

### Figure S7

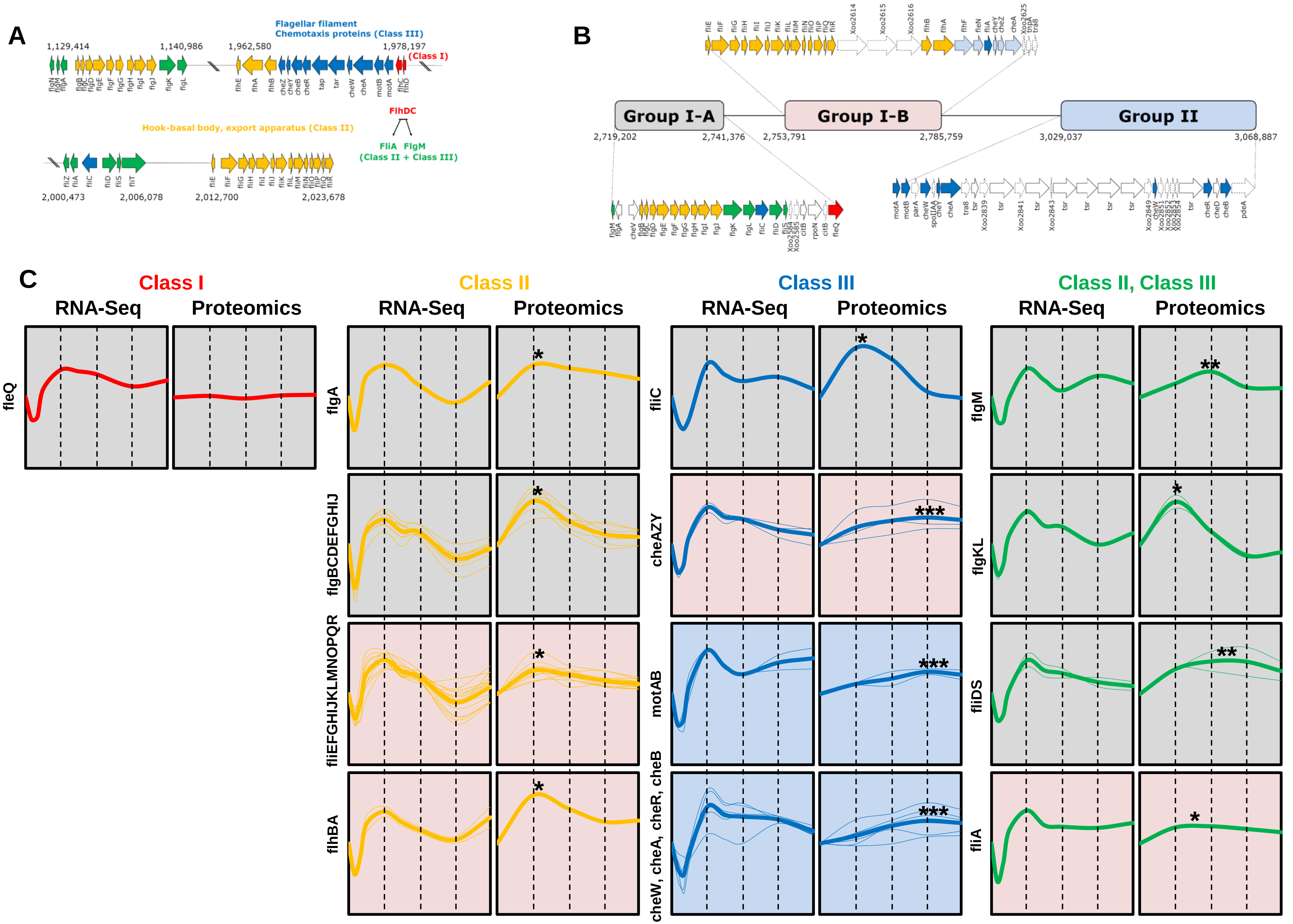

A
Flagellar filament
Chemotaxis proteins (Class III)
(Class I)
Hook-basal body, export apparatus (Class II)
FlhDC
FliA FlgM
(Class II + Class III)
B
Group I-A
Group I-B
Group II
C
Class I
Class II
Class III
Class II, Class III
RNA-Seq
Proteomics
fleQ
flgA
flgBCDEFGHIJ
fliEFGHIJKLMNOPQR
flhBA
fliC
cheAZY
motAB
cheW, cheA, cheR, cheB
flgM
flgKL
fliDS
fliA

### Figure S8

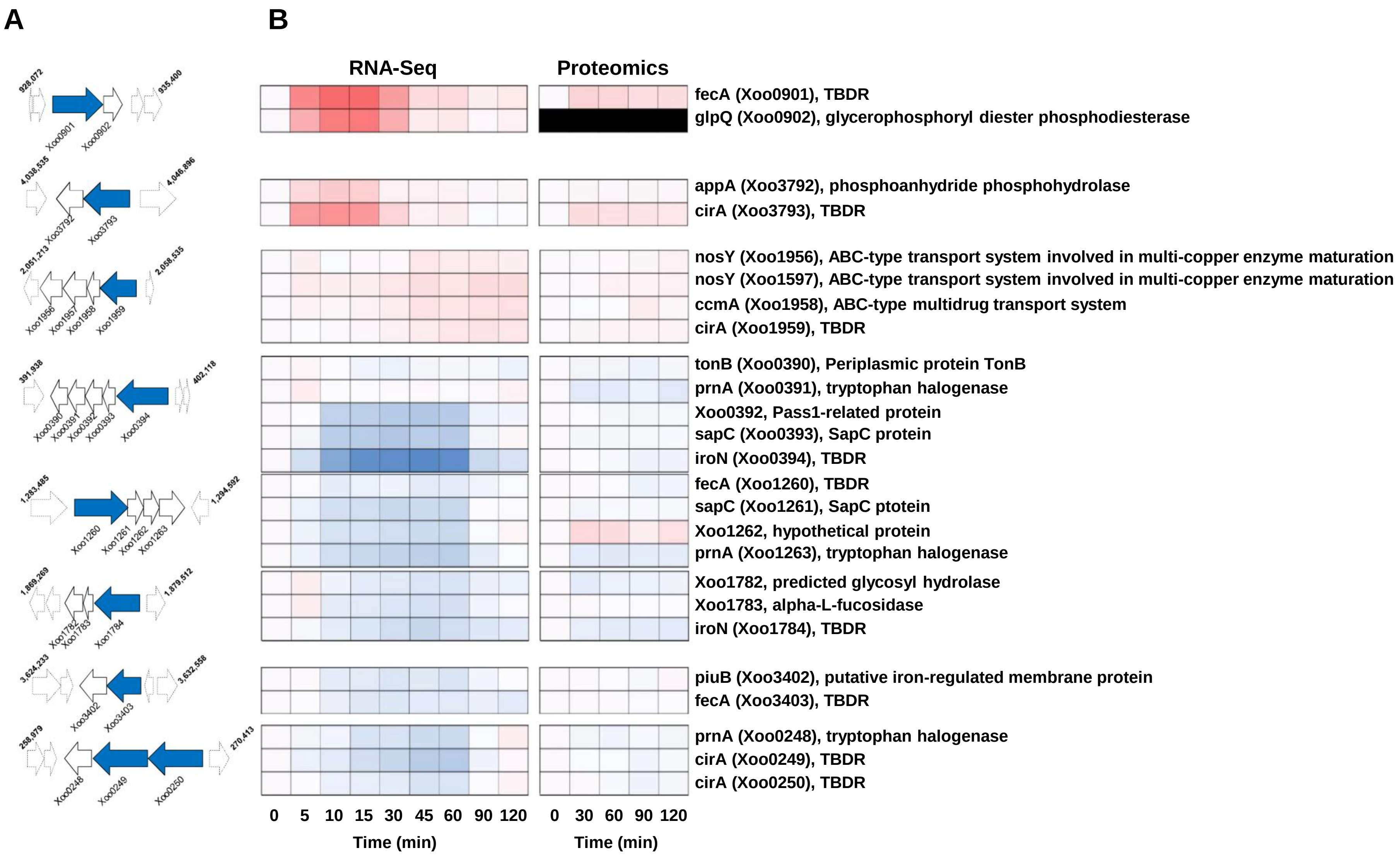
A
B
RNA-Seq
Proteomics
fecA (Xoo0901), TBDR
glpQ (Xoo0902), glycerophosphoryl diester phosphodiesterase
appA (Xoo3792), phosphoanhydride phosphohydrolase
cirA (Xoo3793), TBDR
nosY (Xoo1956), ABC-type transport system involved in multi-copper enzyme maturation
nosY (Xoo1597), ABC-type transport system involved in multi-copper enzyme maturation
ccmA (Xoo1958), ABC-type multidrug transport system
cirA (Xoo1959), TBDR
tonB (Xoo0390), Periplasmic protein TonB
prnA (Xoo0391), tryptophan halogenase
Xoo0392, Pass1-related protein
sapC (Xoo0393), SapC protein
iroN (Xoo0394), TBDR
fecA (Xoo1260), TBDR
sapC (Xoo1261), SapC ptotein
Xoo1262, hypothetical protein
prnA (Xoo1263), tryptophan halogenase
Xoo1782, predicted glycosyl hydrolase
Xoo1783, alpha-L-fucosidase
iroN (Xoo1784), TBDR
piuB (Xoo3402), putative iron-regulated membrane protein
fecA (Xoo3403), TBDR
prnA (Xoo0248), tryptophan halogenase
cirA (Xoo0249), TBDR
cirA (Xoo0250), TBDR
0 5 10 15 30 45 60 90 120
Time (min)
0 30 60 90 120
Time (min)
Xoo0901
Xoo0902
Xoo3792
Xoo3793
Xoo1956
Xoo1957
Xoo1958
Xoo1959
Xoo0390
Xoo0391
Xoo0392
Xoo0393
Xoo0394
Xoo1260
Xoo1261
Xoo1262
Xoo1263
Xoo1782
Xoo1783
Xoo1784
Xoo3402
Xoo3403
Xoo0248
Xoo0249
Xoo0250

### Figure S9

**RNA-Seq** **Proteomics**

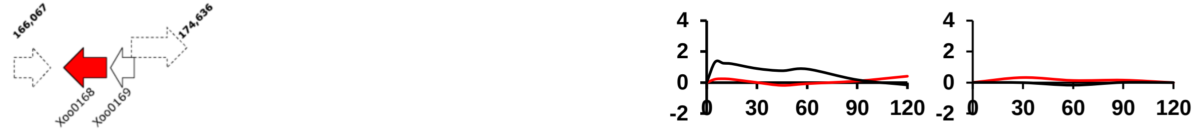


avrBs2 (Xoo0168), avirulence protein
xylR (Xoo0169), xylose repressor-like protein

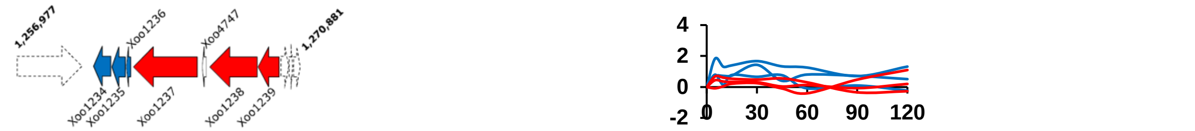


tra8 (Xoo1234), transposase and inactivated derivatives, IS30 family
tra5 (Xoo1235), transposase
Xoo1236, transposase
avrB6 (Xoo1237), avirulence protein
Xoo4747, hypothetical protein
Xoo1238, avirulence protein
avrXa3 (Xoo1239), avirulence protein

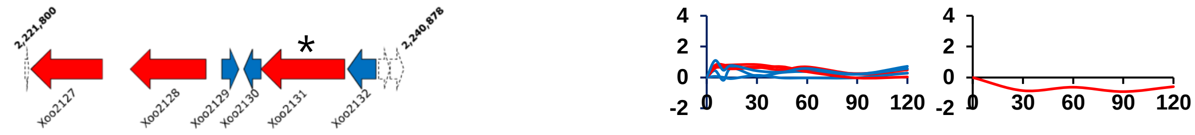


Xoo2127, avirulence protein
Xoo2128, avirulence protein
tra5 (Xoo2129), transposase
Xoo2130, transposase
pthA1 (Xoo2131), avirulence protein
Xoo2132, transposase

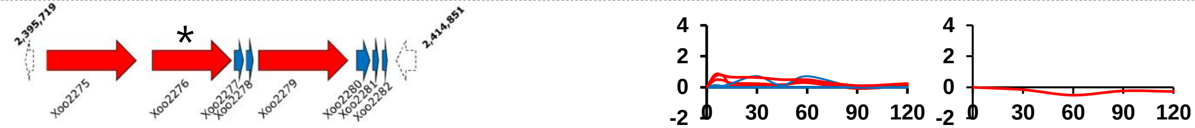


pthA4 (Xoo2275), avirulence protein
Xoo2276, avirulence protein
Xoo2277, transposase
Xoo2278, transposase and inactivated derivatives
Xoo2279, avirulence protein
tra8 (Xoo2280), transposase
tra8 (Xoo2281), transposase
tra8 (Xoo2282), transposase

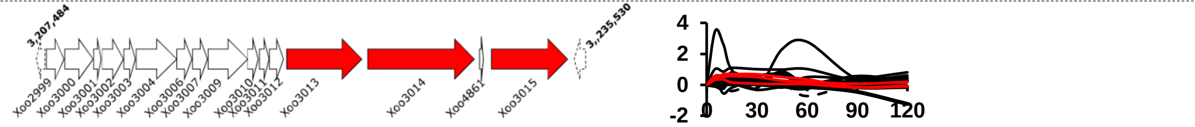


cybB (Xoo2999), cytochrome B561
Xoo3000, cytochrome C
cycA (Xoo3001), cytochrome C2
Xoo3002, hypothetical protein
rpoE (Xoo3003), RNA polymerase sigma-E factor
aprE (Xoo3004), subtilisin-like serine protease
ccmC (Xoo3006), cytochrome C-type biogenesis protein
cycJ (Xoo3007), cytochrome C-type biogenesis protein
cycK (Xoo3009), cytochrome C-type biogenesis membrane protein
dsbE (Xoo3010), cytochrome C-type biogenesis protein/thioredoxin
cycL (Xoo3011), cytochrome C-type biogenesis protein
Xoo3012, cytochrome C biogenesis factor
pthA2 (Xoo3013), avirulence protein
Xoo3014, avirulence protein
Xoo4861, hypothetical protein
pthA3 (Xoo3015), avirulence protein

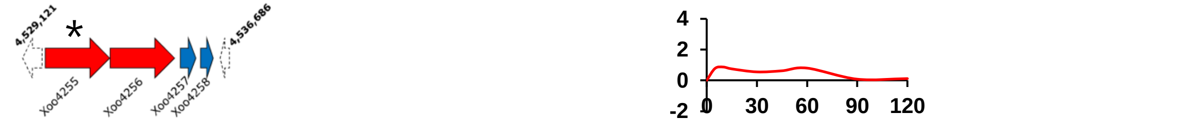


avrXa7 (Xoo4255), avirulence/virulence protein
avrXa3 (Xoo4256), avirulence protein
Xoo4257, transposase
Xoo4258, transposase
